# Fibroblast alignment coordinates epithelial migration and maintains intestinal tissue integrity

**DOI:** 10.1101/2021.05.28.446131

**Authors:** Jordi Comelles, Aina Abad-Lázaro, Verónica Acevedo, David Bartolomé-Català, Aitor Otero-Tarrazón, Anna Esteve-Codina, Xavier Hernando-Momblona, Eduard Batlle, Vanesa Fernández-Majada, Elena Martinez

## Abstract

Fibroblasts reside underneath most epithelial tissues. In the intestine, recent studies have shown that fibroblast migration contributes to tissue morphogenesis and wound healing. Yet, whether physical interactions between epithelial cells and fibroblasts contribute to epithelial movement remains elusive. Here, we show that subepithelial fibroblast alignment enhances directed and persistent migration of organoid-derived intestinal epithelia. Using a reconstituted epithelial–stromal gap-closure model, we demonstrate that direct contact with fibroblasts improves gap closure by promoting cell alignment, sustaining tissue integrity, and synchronizing crypt–villus migration. Fibroblasts undergo long-range ordering to align perpendicularly to the epithelial front and deposit protein paths that act as guidance features to direct epithelial migration. In parallel, epithelial cells acquire a wound-associated epithelial-like phenotype, but insufficient to explain the effects of fibroblast contact. Our findings uncover a dual role for intestinal fibroblasts in epithelial repair, coordinating both biochemical and physical cues to ensure efficient and cohesive migration.

## Introduction

Fibroblasts reside underneath epithelial layers in most organs. They constitute the primary component of a supportive mesenchymal compartment that interfaces with epithelial cells via a basement membrane primarily composed of laminin and type IV collagen.^1^ Beyond providing physical support, subepithelial fibroblasts play a crucial role in regulating various epithelial processes throughout development, homeostasis, and disease. For instance, they restrain duct elongation and control branching in mammary glands,^2^ and regulate bronchial epithelial repair during lung inflammation or injury.^3^ Also, analyses carried out in skin,^4^ cornea,^5,6^ or cleaved palate,^7^ indicate that fibroblasts contribute to epithelial restoration upon injury.^8,9^ Thus, understanding the complex interplay between fibroblasts and epithelia across different contexts and how they mechanistically integrate for tissue function, is a fundamental question. This has broad implications for tissue engineering, developmental biology, and regenerative medicine, offering insights into both physiological maintenance and pathological conditions.

In the intestinal tract, the epithelial layer is also supported by fibroblasts present in the lamina propria.^10–13^ The different subpopulations of these intestinal fibroblasts vary in morphology, secretory profile and localization along the crypt-villus axis that compartmentalizes the intestinal epithelium.^14^ They are known to play a crucial role in maintaining epithelial homeostasis by secreting various factors^15,16^ including Wnts,^17^ R-spondins,^16,18^ Bone Morphogenetic Protein (BMP) agonists and antagonists,^14,19–22^ as well as Epidermal Growth Factor Receptor (EGFR) ligands.^23^ However, their role is not solely restricted to provide paracrine signaling but they also physically interact with epithelial cells. For example, migration and aggregation of Platelet Derived Growth Factor Receptor A (PDGFRA)^+^ subepithelial fibroblasts seems sufficient for generating the curvature needed to drive villi formation during intestinal development in mice.^24^

Moreover, in vivo wound healing experiments in colonic epithelium have underscored the significance of both the epithelial and mesenchymal components in tissue repair.^25–28^ The epithelial integrity hinges on the migration of non-proliferative wound-associated epithelial (WAE) cells, originating from crypts adjacent to the wound and moving over the wound surface.^25^ The differentiation of these specialized WAE cells is regulated by the signaling of fibroblasts localized at the site of injury producing Prostaglandin E_2_ (PGE_2_).^27,28^ However, there is still limited understanding regarding the early migration of intestinal epithelial cells following injury. In particular, it remains unclear whether and how the physical interaction between these cells and intestinal fibroblasts influences epithelial migration and function restoration beyond fibroblasts’ secretory function.

Early in vitro studies have shown that intestinal epithelial Caco-2 cells close wounds more efficiently when cultured on intestinal myofibroblasts compared to when cultured on extracellular matrix (ECM)-coated plastic plates.^29^ However, this cancer-derived cell line does not accurately replicate the unique migratory dynamics of the in vivo intestine,^30^ and the mechanisms behind fibroblast-mediated migration are not well understood. Recently, intestinal and colon organoids have been successfully opened-up into 2D monolayers.^31,32^ These monolayers retrieve their crypt-and villus-like organization, facilitating the study of signaling feedback loops and mechanical patterning and migration.^31,33,34^ Despite this progress, the role of the stromal compartment has been largely overlooked. Here, using a bioengineered epithelial–stromal model comprising organoid-derived intestinal epithelial cells (IECs), primary intestinal fibroblasts, and a basement membrane matrix, we provide evidence that the physical interaction between intestinal fibroblasts and epithelia induces a long-range organization of fibroblasts, which, in its turn, results in a directed and persistent migration of epithelial cells leading to the restoration of intestinal epithelial tissue in an in vitro gap closure model.

## Results

### Fibroblast contact enhances organoid-derived intestinal epithelium gap closure in vitro

To investigate the role of the physical crosstalk between IECs and intestinal fibroblasts, we set up a physiological-like model of intestinal mucosa in vitro (Figure 1a). This model included primary mouse intestinal subepithelial fibroblasts and organoid-derived intestinal mouse epithelial cells, which grew as self-organized monolayers following an in-house developed protocol.^32,35^

**Figure 1.**
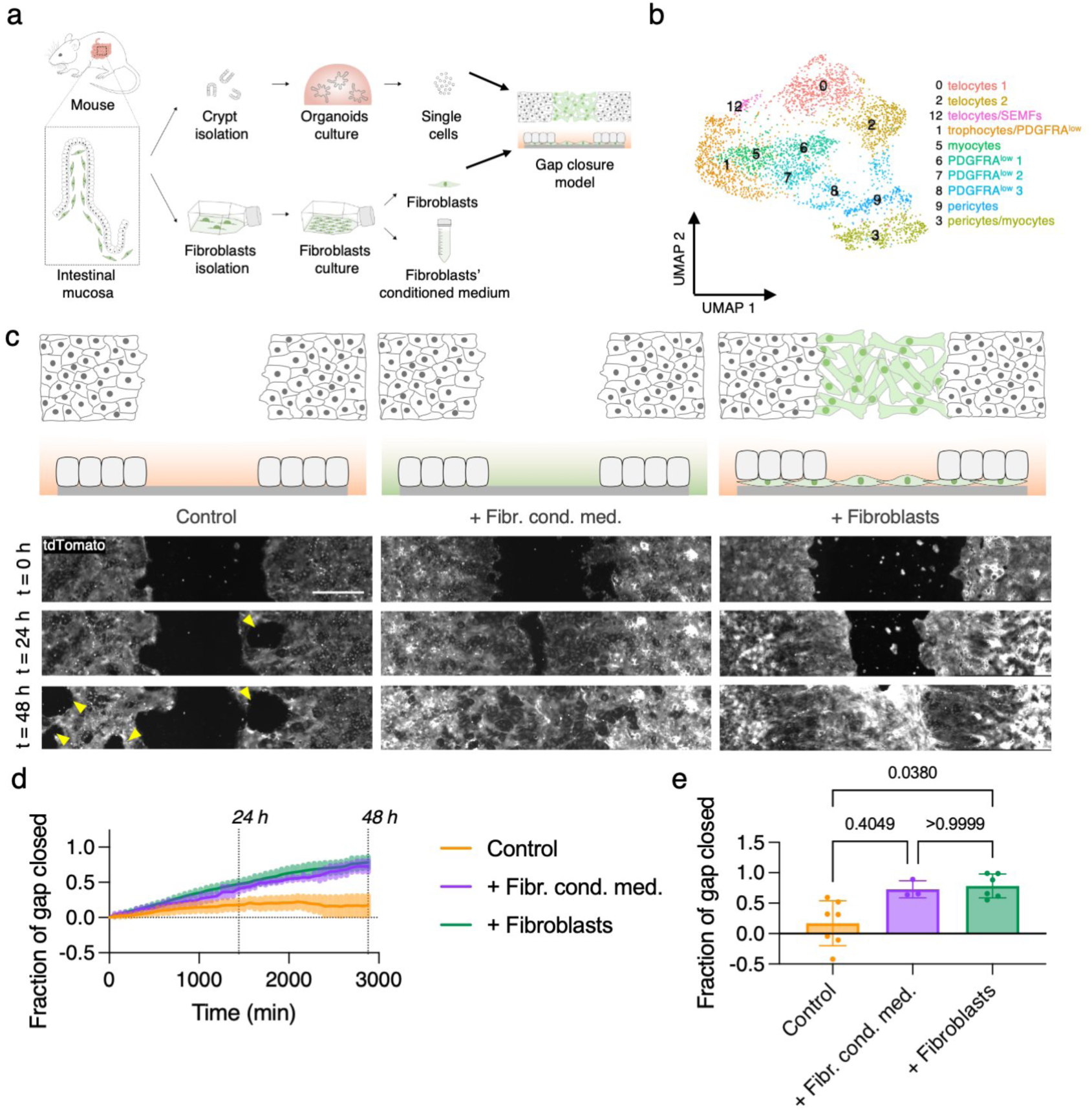
Primary intestinal fibroblasts promote epithelial gap closure: (a) Schematics of the experimental set-up. (b) Uniform manifold approximation and projection (UMAP) of established primary culture from small intestine stroma. Resident populations were dominated by molecularly distinct clusters of telocytes, pericytes, myocytes and PDGFRA^low^ cells. (c) Schematics of the experimental set-up (top). Snapshots of the live-imaging of tdTomato organoid-derived cells migrating in control, + fibr. cond. med. and + fibroblasts conditions at 0, 24 and 48h upon barrier removal (bottom). Yellow arrows highlight holes in the monolayers. Scale bar: 500 µm. (d) Fraction of gap closed along time in control, + fibr. cond. med. and + fibroblasts conditions. Mean ± SEM. (e) Fraction of gap closed at 48 in control, + fibr. cond. med. and + fibroblasts conditions. Mean ± SD. N = 7, N = 3 and N = 6 independent experiments for control, + fibr. cond. med. And + fibroblasts conditions, respectively. Statistical significance was assessed using a Kruskal-Wallis test.

Primary fibroblasts were isolated from mouse intestinal mucosa and cultured on standard tissue culture plates. Cells were then stained for mesenchymal markers (α-SMA, vimentin, desmin, FOXL1, PDGFRα) and ECM proteins (fibronectin, laminin, and collagen IV). The majority of the cells were vimentin-positive (Figure S1a), and a substantial fraction also expressed α-SMA (Figure S1b). Additionally, subsets of cells were positive for FOXL1 (Figure S1c) and PDGFRα (Figure S1d), while only a few expressed desmin (Figure S1e). These fibroblasts also expressed and secreted key ECM proteins, including fibronectin (Figure S1a), laminin (Figure S1b), and collagen IV (Figure S1e). This marker profile suggests the presence of diverse subepithelial mesenchymal populations.^14–16^

To explore this diverse cellular composition of the isolated population, we performed single-cell RNA sequencing (scRNA-seq) [see Supplementary Material and Figure S2]. A small subset of cells formed a *Pecam1*⁺ cluster corresponding to endothelial cells, while another *Gfap*⁺ cluster was assigned to neurons and glial cells. Additionally, two clusters expressing *Cd52* at varying levels were identified as immune cells. These non-fibroblastic cell types, commonly found in primary intestinal stromal preparations, accounted for less than 15% of the total cell population.^14^ The remaining ∼85% of cells, comprising the dominant population, were grouped into 10 transcriptionally distinct fibroblast subclusters (Figure 1b). Although marker expression was not strictly restricted to a single subtype, analysis of canonical and tissue-specific markers revealed distinct clusters corresponding to known stromal cell types, including pericytes, telocytes/subepithelial myofibroblasts (SEMFs), trophocytes, myocytes, and PDGFRA^low^ stromal cells. These subpopulations closely mirrored the transcriptional profiles of fibroblastic cell types previously described in proximity to the intestinal epithelium.^14^ Notably, many of these fibroblasts expressed key ligands and modulators of the Wnt and BMP signaling pathways, suggesting their potential roles in supporting epithelial cell proliferation and differentiation.^17,22,36^

Next, to study epithelial restoration we grew a monolayer of these intestinal fibroblasts on a thin layer of Matrigel. Then, we placed customized elastomeric barriers on top of this monolayer. Subsequently, we seeded IECs derived from organoids on top the fibroblasts. After the formation of the epithelial monolayer, we removed the elastomeric barrier creating an epithelium-free gap, and we proceeded to observe and track for 48 hours the migration of epithelial cells attempting to close the gap (+ fibroblasts condition). To unravel the potential role of fibroblasts in this gap closure process, we tested two additional gap conditions: (i) epithelial cells alone (control), and (ii) epithelial cells cultured with fibroblasts’ conditioned medium (+ fibr. cond. med.) (Figure 1c).

At the moment of the removal of the elastomeric barrier, confluent epithelial monolayers were present on both sides of the gap across all conditions, regardless of whether fibroblasts or their conditioned medium were present (Figure 1c, t = 0 hours, Figure S3a-c). In addition, the intestinal epithelial monolayers exhibited their characteristic crypt-like and villus-like compartmentalization, as previously described.^31,32,34^ This organization was marked by densely packed EphB2⁺ regions containing proliferative (Ki-67⁺) cells, along with Paneth cells (Lysozyme⁺/Wnt3a⁺) and adjacent Wnt3a⁺ stem-like cells, consistent with the crypt compartment (Figure S3d–f). Surrounding these regions were non-proliferative (Ki-67⁻), EphB2⁻, Fabp1⁺ and/or Ck20⁺ cells, indicative of villus-like differentiation (Figure S3d–f).

After 24 hours, noticeable differences emerged among the conditions. In the + fibr. cond. med. and + fibroblasts conditions, the epithelial front advanced, resulting in significant gap closure. In contrast, the controls exhibited poor epithelial movement and a disruption in the epithelial cohesion, as seen by the emergence of holes within the monolayers (Movie 1 and Figure 1c, t = 24 hours, indicated by yellow arrow). After 48 hours, gap closure failed for the control monolayers, and the number and size of holes within the tissue had significantly increased (Figure 1c, t = 48 hours, indicated by yellow arrows). Meanwhile, the + fibr. cond. med. and + fibroblasts conditions succeeded in closing the gap (Figure 1d). Actually, epithelial monolayers achieved the most effective gap closure when cultured in direct physical contact with the fibroblasts (Figure 1e and Movies 2 and 3).

### Fibroblast-enhanced gap closure is independent of PGE₂–EP4 signaling

Traditionally, intestinal epithelial migration has been attributed to mitotic pressure generated by cell proliferation within the crypts.^37,38^ As cells migrate upward along the villus axis, they continue to move actively even in the absence of proliferation, undergoing differentiation along the way.^30^ However, during in vivo wound healing, gap closure follows a different mechanism. In this context, stromal-derived PGE₂ signals through the EP4 receptor to drive intestinal stem cells, progenitor cells, and likely immature enterocytes to differentiate into WAE cells. This strategy enables the epithelium to close the wound efficiently without relying on an increase of proliferation.^25,28^

Given this, we next asked whether differences in cell proliferation or activation of the WAE cell program could explain the distinct gap closure behaviors observed between control epithelial monolayers and those migrating in the presence of stromal fibroblasts. To assess the contribution of proliferation, we quantified Ki-67⁺ cells both within the epithelial monolayer and at the leading edge of migration (Figure 2a and Figure S4a,b). Strikingly, all three experimental conditions exhibited similar patterns, with no significant differences between them. Within the monolayers, Ki-67^+^ cells accounted for approximately 5% of the population (Figure 2b), typically forming clusters associated with crypt-like domains.^31,32,34^ In contrast, the migration front (first 250 µm of the epithelial front) consistently showed a higher proportion of proliferative cells, exceeding 20% across all conditions (Figure 2b). Similarly, staining for Cleaved Caspase-3 (CC3) revealed low and comparable levels of apoptosis across the three conditions (Figure 2a and Figure S4a,b). These results suggest that while proliferation and survival may contribute to epithelial migration, they do not account for the differences in gap closure efficiency observed between conditions.

**Figure 2.**
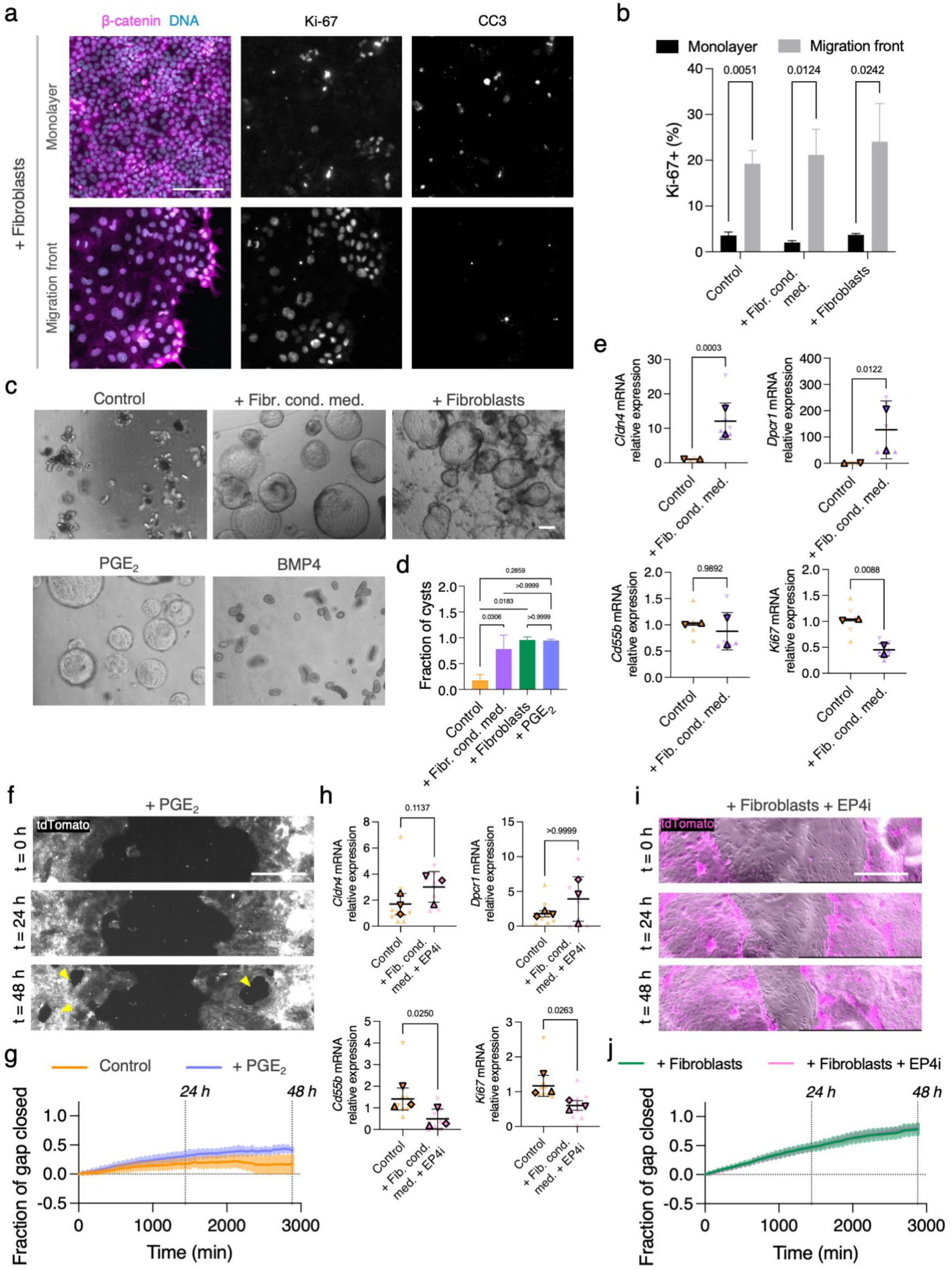
Fibroblasts induce a WAE-like program via PGE₂, but promote gap closure independently. (a) Representative sum intensity projections images of epithelial cells in the monolayer region and in the migration front region at 24h upon barrier removal immunostained for DAPI (DNA), β-Catenin, Ki-67 and CC3 of + fibroblasts condition. Scale bar: 100 µm. (b) Percentage of Ki-67^+^ cells in the monolayer and in the migration front region of control, + fibr. cond. med. and + fibroblasts conditions. Mean ± SEM. Two-way ANOVA test. (c) Representative bright field images of crypts grown in 3D Matrigel® drops with ENRCV medium (control), with fibroblasts conditioned medium (+ fibr. cond. med.), in coculture with primary fibroblasts (+ fibroblasts), with ENRCV medium supplemented with PGE_2_ and supplemented with BMP4 after 2 days in culture. Scale bar: 1000 µm. (d) Fraction of cystic organoids respect to all organoids per condition. Mean ± SD. Kruskal-Wallis test. (e) *Cldn4*, *Dpcr1*, *Cd55b* and *Ki67* mRNA relative expression for control and + fibr. cond. Med 3D Matrigel® drops. Mean ± SD. N = 6 drops from 2 independents experiments per condition. Statistical significance was assessed using a Kruskal-Wallis test. (f) Snapshots of the live-imaging of mT/mG organoid-derived cells migrating in PGE_2_ conditions at 0, 24 and 48h upon barrier removal. Yellow arrows highlight holes in the monolayers. Scale bar: 500 µm. (g) Fraction of gap closed along time in control and + PGE_2_ conditions. Mean ± SEM. N = 7 and N = 3 independent experiments respectively. (h) *Cldn4*, *Dpcr1*, *Cd55b* and *Ki67* mRNA relative expression for control and + fibr. cond. med. + EP4i 3D Matrigel® drops. Mean ± SD. N = 9 drops from 3 independents experiments per condition. Statistical significance was assessed using a Kruskal-Wallis test. (i) Snapshots of the live-imaging of mT/mG organoid-derived cells and fibroblasts migrating in +fibroblasts + EP4i conditions at 0, 24 and 48h upon barrier removal. Scale bar: 500 µm. (j) Fraction of gap closed along time in + fibroblasts and +fibroblasts + EP4i conditions. Mean ± SEM. N = 6 and N = 4 independent experiments respectively.

To further explore the role of PGE₂ signaling and WAE cell induction, we conducted 3D co-culture experiments in Matrigel. Intestinal crypts were embedded either alone, with intestinal fibroblasts, with their conditioned medium, or in media supplemented with PGE₂ or BMP4 (Figure 2c), which are reported to be important fibroblasts signals.^28,39^ Over a span of four days, we monitored crypt development and morphological changes. By day two, control crypts developed into typical organoids with budding crypt-like structures and interspersed villus-like regions, reflecting normal epithelial organization. In contrast, when cultured with fibroblasts or their conditioned medium, crypts developed into cystic structures lacking budding domains (Figure 2c,d). Notably, this cystic morphology was also observed in crypts treated with PGE₂, but not with BMP4, suggesting that PGE₂ signaling may be present among fibroblast-derived factors.

Although cyst formation has previously been linked to increased proliferation,^16–18,40^ we did not observe a higher proportion of Ki-67⁺ cells in the conditioned medium condition compared to controls (Figure S4c). Rather, the spatial distribution of proliferative cells was altered: while Ki-67⁺ cells in control organoids were confined to crypt-like domains, they appeared randomly distributed in cysts formed in the presence of conditioned medium.

Given these morphological similarities between crypts cultured with fibroblasts, conditioned medium, and PGE₂, we next asked whether these epithelial cells acquired a WAE-like identity. Using quantitative reverse transcription polymerase chain reaction (RT-qPCR), we measured the expression of WAE–associated genes.^28^ Crypts exposed to fibroblast-conditioned medium showed significantly increased expression of *Cldn4* and *Dpcr1*, while *Cd55b* remained unchanged (Figure 2e). Additionally, these cells expressed lower levels of *Ki67*, the stem cell marker *Lgr5*, and the differentiation marker *Fabp1*, together with lower levels of *Axin2*, a transcriptional readout for canonical Wnt signaling (Figure 2e and Figure S4d). This transcriptional profile is consistent with a less proliferative, less differentiated, WAE-like state, similar to that observed during in vivo wound repair.

To test whether PGE₂ alone could promote epithelial migration, we added it to control cultures. Although gap closure improved compared to untreated controls, the epithelial monolayer still failed to fully close the gap, with holes forming during migration, similar to control conditions (Figure 2f and Movie 4). Quantification confirmed this partial improvement: PGE₂-treated monolayers closed a larger fraction of the gap than controls but less than those co-cultured with fibroblasts (Figure 2g and Figure S4e).

To directly assess the role of PGE₂–EP4 signaling, we inhibited the EP4 receptor in epithelial monolayers migrating on fibroblasts. Inhibition reversed the WAE-like transcriptional program, reducing *Cldn4*, *Dpcr1*, and *Cd55b* expression, while restoring *Lgr5* and *Axin2* to control levels (Figure 2h and Figure S4f). Surprisingly, however, EP4 inhibition did not impair migration. Epithelial cells still closed the gap efficiently, reaching levels comparable to untreated fibroblast co-cultures (Figure 2i,j, Figure S4e and Movie 5). Moreover, *Ki67* and *Fabp1* expression remained low even after EP4 inhibition (Figure S4f), suggesting that fibroblast-induced changes in proliferation and differentiation extend beyond the PGE₂–EP4 axis.

Taken together, these findings indicate that while fibroblast-derived signals promote a WAE-like gene expression profile, the enhanced efficiency of epithelial migration and gap closure is independent of PGE₂–EP4 signaling. Additionally, live imaging revealed that fibroblasts physically interact with the epithelium. On one hand when holes formed in the monolayer, fibroblasts were frequently observed at those sites, where they appeared to facilitate rapid epithelial re-spreading and sealing (Movie 6). On the other hand, the coculture in Matrigel drops, led to the expansion of the epithelium over the surface, leading to the transformation of cysts into flat monolayers exhibiting the characteristic hexagonal packing pattern of epithelial cells by day 14 of coculture (Figure S5a, b). We observed that fibroblasts were arranged perpendicular to the cysts and were physically interacting with them (Figures S5a, c, d and Movie 7), suggesting a potential pulling of the fibroblasts on the epithelial cells. This suggests a mechanical or contact-mediated role for fibroblasts in preserving epithelial integrity and promoting coordinated migration besides their paracrine signaling.

### Fibroblasts coordinate crypt–villus migration and preserve epithelial integrity

To understand how fibroblasts promote efficient gap closure, we first examined the migratory behavior of crypt and villus epithelial cells. In control conditions, we observed a striking dissociation between crypt and villus cell dynamics: while villus epithelial cells migrated toward the gap, crypt cells remained largely static, resulting in mechanical uncoupling between these functionally distinct regions (Figure 3a and Movie 8). This lack of coordination persisted when epithelia were exposed to fibroblast-conditioned medium, indicating that soluble signals alone were insufficient to synchronize tissue-scale movement. Only the physical presence of underlying fibroblasts fully synchronized crypt-villus dynamics, significantly increasing both the net displacement and total migratory distance of crypt cells (Figure 3b,c and Movie 9).

**Figure 3.**
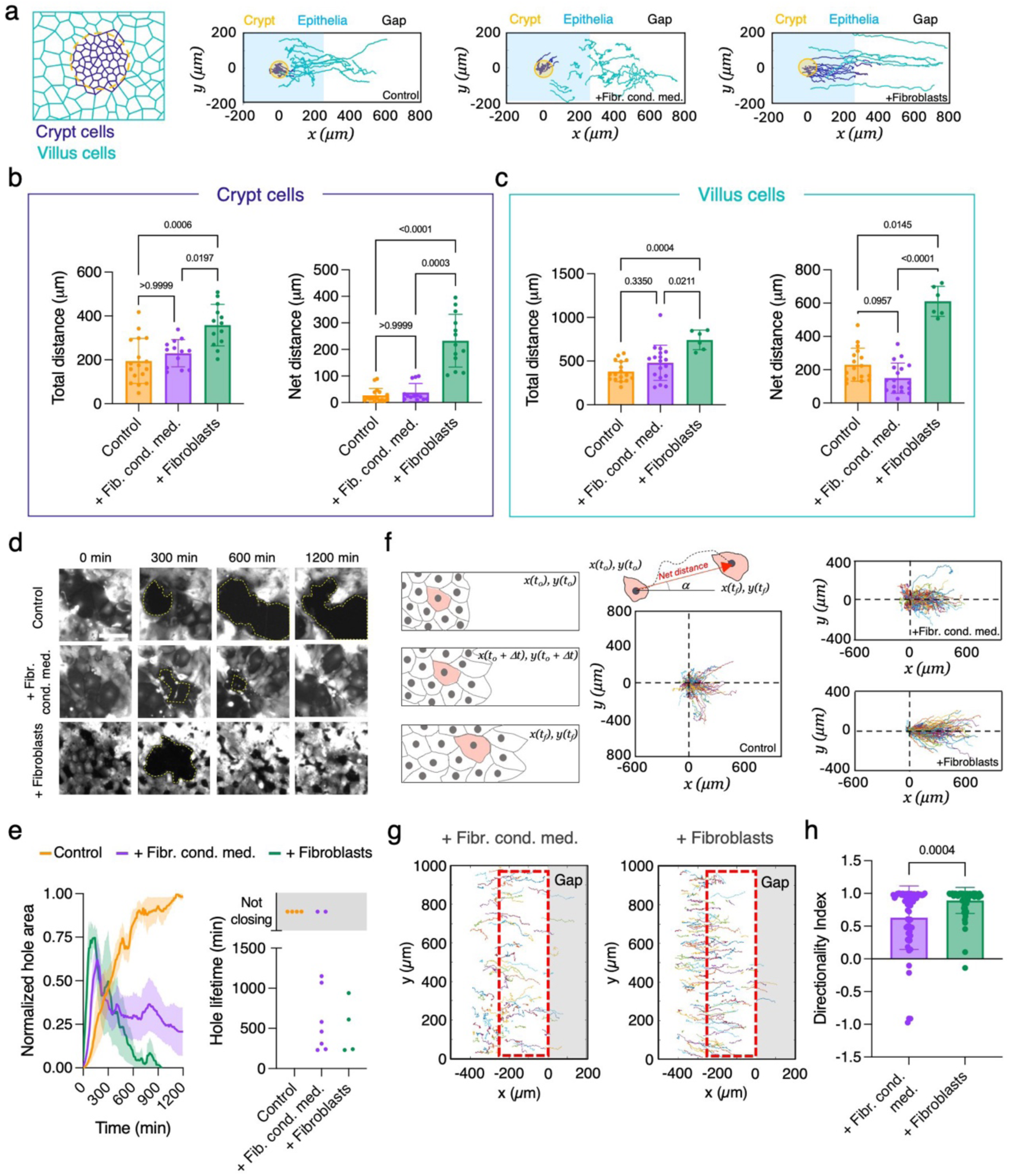
Physical contact with fibroblasts enhances directional epithelial migration and preserves integrity by coordinating crypt–villus dynamics. (a) Trajectories of individual cells from the crypt and from the villus regions in control, + fibr. cond. med. and + fibroblasts conditions. The direction of the gap is at x > 0. (b) Total distance and net distance covered by crypt cells in control, + fibr. cond. med. and + fibroblasts conditions. Mean ± SD and individual values. Kruskal-Wallis test. N= 5, N = 3 and N = 3 independent experiments respectively. (c) Total distance and net distance covered by villus cells in control, + fibr. cond. med. and + fibroblasts conditions. Mean ± SD and individual values. Kruskal-Wallis test. N= 4, N = 4 and N = 2 independent experiments respectively. (d) Snapshots of the live-imaging of tdTomato organoid-derived cells migrating in control, + fibr. cond. med. and + fibroblasts conditions. t=0 min corresponds to right before hole emergence. Holes areas are delineated with a dotted white line. Scale bar: 100 µm. (e) Left: Time-dependent change in hole area normalized by the maximum area of that hole in control, + fibr. cond. med. and + fibroblasts conditions. Mean ± SEM. Right: Lifetime of holes in control, + fibr. cond. med. and + fibroblasts conditions. N = 4, N = 9 and N = 4 holes respectively. (f) Schematics of the tracking analysis and the net displacement vector. Trajectories of each individual cell centered at the origin at t = 0. The direction of the gap is at x > 0. (g) Cell trajectories for + fibr. cond. med. and + fibroblasts conditions. The gray region corresponds to the gap while the red rectangle highlights the first 250 µm of the epithelium behind the migration front. (h) Directionality index for cells’ trajectories within the first 250 µm behind the migration front. Mean ± SD and individual values. Kruskal-Wallis test.

We next asked how this coordination influenced epithelial integrity. In control monolayers, we frequently observed intra-epithelial hole formation, particularly in regions adjacent to large, likely differentiated cells. Time-lapse analysis revealed that holes initiated with rupture of cell-cell contacts, followed by cell retraction and failed closure attempts by neighbors (Figure 3d,e and Movie 10). The spatial pattern of hole formation suggested that mechanical stresses generated by uncoordinated crypt-villus movement may contribute to epithelial fragility. In contrast, monolayers cultured in the physical presence of fibroblasts showed dramatically different behavior. While holes still formed, they were rapidly resolved, with epithelial integrity restored within hours (Figure 3d,e and Movie 11). Fibroblast-conditioned medium led to an intermediate phenotype, enabling hole closure but with reduced efficiency compared to direct contact conditions (Figure 3d,e and Movie 12).

To quantify how fibroblasts enhanced directed migration and tissue cohesion, we analyzed single-cell trajectories across the monolayer (Figure 3f and Figure S6a-c). Epithelial cells migrating on fibroblasts exhibited highly directional movement toward the gap, with alignment values (cos(2α)) approaching 1 within 250 µm of the front (Figure 3g, h and Figure S6d). In contrast, control trajectories showed no such alignment (Figure 3f and Figure S6d), while fibroblast-conditioned medium induced partial improvement (Figure 3g, h and Figure S6d).

Together, these findings demonstrate that fibroblasts promote gap closure through a dual mechanism: enhancing the persistence of individual epithelial migration and synchronizing movement across crypt and villus compartments. The physical presence of fibroblasts appears critical for this coordination, likely through direct epithelial–stromal contact or matrix-mediated signaling, providing the spatial and mechanical cues needed to couple epithelial migration with tissue integrity.

### Fibroblasts migrate toward the epithelial front and become activated during gap closure

Results above point toward the existence of physical interactions between the IECs and the fibroblasts that collaborate to regulate epithelial migration. To better understand these interactions, we studied more in detail the gap closure process in the + fibroblasts condition. First, we stained the gap region at various time points to examine the epithelial front and the distribution of fibroblasts close to it (Figure 4a). Upon removal of the barrier (t = 0 h), fibroblasts at the epithelial front were small and evenly distributed, with no prominent α-SMA fibers present. However, 24 and 48 hours after barrier removal, fibroblasts were more abundant and appeared bigger and activated, exhibiting prominent α-SMA fibers, both beneath the epithelial front and within the gap region closer to the leading edge. Then, we analyzed the onset of gap closure (first 9 hours after removing the barrier) using particle image velocimetry (PIV), which allowed us to obtain velocity fields from both the IECs and the fibroblasts. Strikingly, two clearly differentiated regions appeared on the maps of the x-component of the cells’ mean velocities (v_x_) (Figure 4b). IECs exhibited on average v_x_ > 0, so they were migrating toward the gap. In contrast, fibroblasts displayed on average v_x_ < 0, indicating that they were migrating from the gap toward the epithelium. The temporal evolution of the x-component of the instantaneous velocity revealed the expansion of the v_x_ > 0 region toward the gap, which correlated with the advancing epithelium (Figure 4c). Furthermore, the averaged velocity profiles of the x-component (v_x_) and the y-component (v_y_) across the migration fronts and adjacent gap regions indicated that migration occurred mainly in the direction perpendicular to the front for both cell types and that fibroblasts in the gap region actively moved toward the front of the migrating epithelium (v_x_ < 0, v_y_ ∼0) (Figure 4d). Altogether, upon barrier removal fibroblasts migrate from the gap towards the leading edge of the epithelia, where they express prominent α-SMA fibers and seem to interact with the epithelial cells. This indicates that not only fibroblasts from underneath the epithelia but also those from further away inside the gap are involved in the gap closure process.

**Figure 4.**
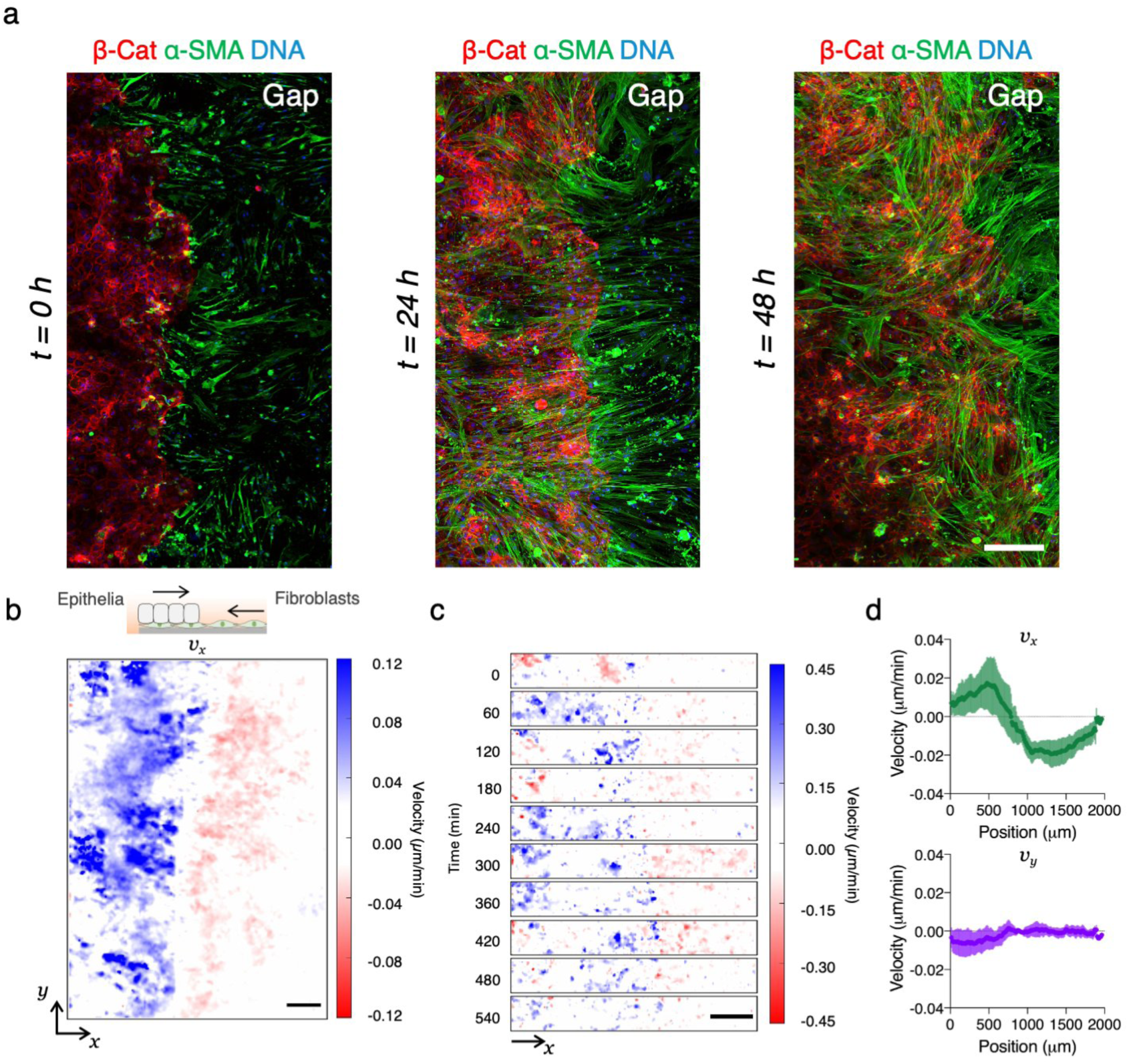
Fibroblasts migrate toward the epithelial front and become activated during gap closure. (a) Representative fluorescence microscopy images corresponding to immunostaining of β-Catenin (β-Cat), alpha-smooth muscle actin (ɑ-SMA) and DNA of + fibroblasts condition. Timeframes corresponding to 0, 24 and 48h upon barrier removal. Scale bar: 100 µm. (b) Mean velocity in the x-component (v_x_) during the first 9 hours of gap closure. Scale bar: 250 µm. (c) Snapshots of the instantaneous velocity v_x_ of a selected region along the first 9 hours of gap closure. Scale bar: 250 µm. (d) Profiles of the x-component (v_x_) and y-component (v_y_) of the mean velocity across the gap direction during the first 9 hours of gap closure. Mean ± SD of n = 4 migration fronts of N = 2 independent experiments.

### Long-range fibroblast alignment leads to spatiotemporal co-orientation with epithelial cells

These results suggest a coordinated and interactive migration process within the gap involving both IECs and the intestinal fibroblasts. So, to gain a deeper understanding of fibroblast movements during epithelial migration we decided to follow the dynamics of fibroblast rearrangement during epithelial migration (Figure 5a). Using primary fibroblasts expressing GFP at the cell membrane,^41^ we examined their arrangement relative to the migration front and analyzed their orientation in the central region of the gap (Figure 5b). At the beginning of the experiment, we observed small swirly patterns in the orientation fields (Figure 5b), indicative of short-range alignments of polarity. However, over time, we observed a progressive alignment of fibroblasts perpendicular to the migration front (orientation vectors around 0°) (Figure 5b,c), resulting in the enlargement of the initial short-range alignments of polarity (Figure 5b). To visualize local discontinuities in alignment, we calculated the nematic order for each vector grid.^42^ Over time, discontinuities between swirly patterns disappeared, indicating long-range alignments of polarity (Figure 5b), and the fraction of disordered regions (nematic order below 0.5)^43^ decreased (Figure 5d). Notably, fibroblasts far away from the migration front (up to 750 µm) were also aligned, strongly suggesting a mechanism of fibroblast-fibroblast communication that results in a long-range order.

**Figure 5.**
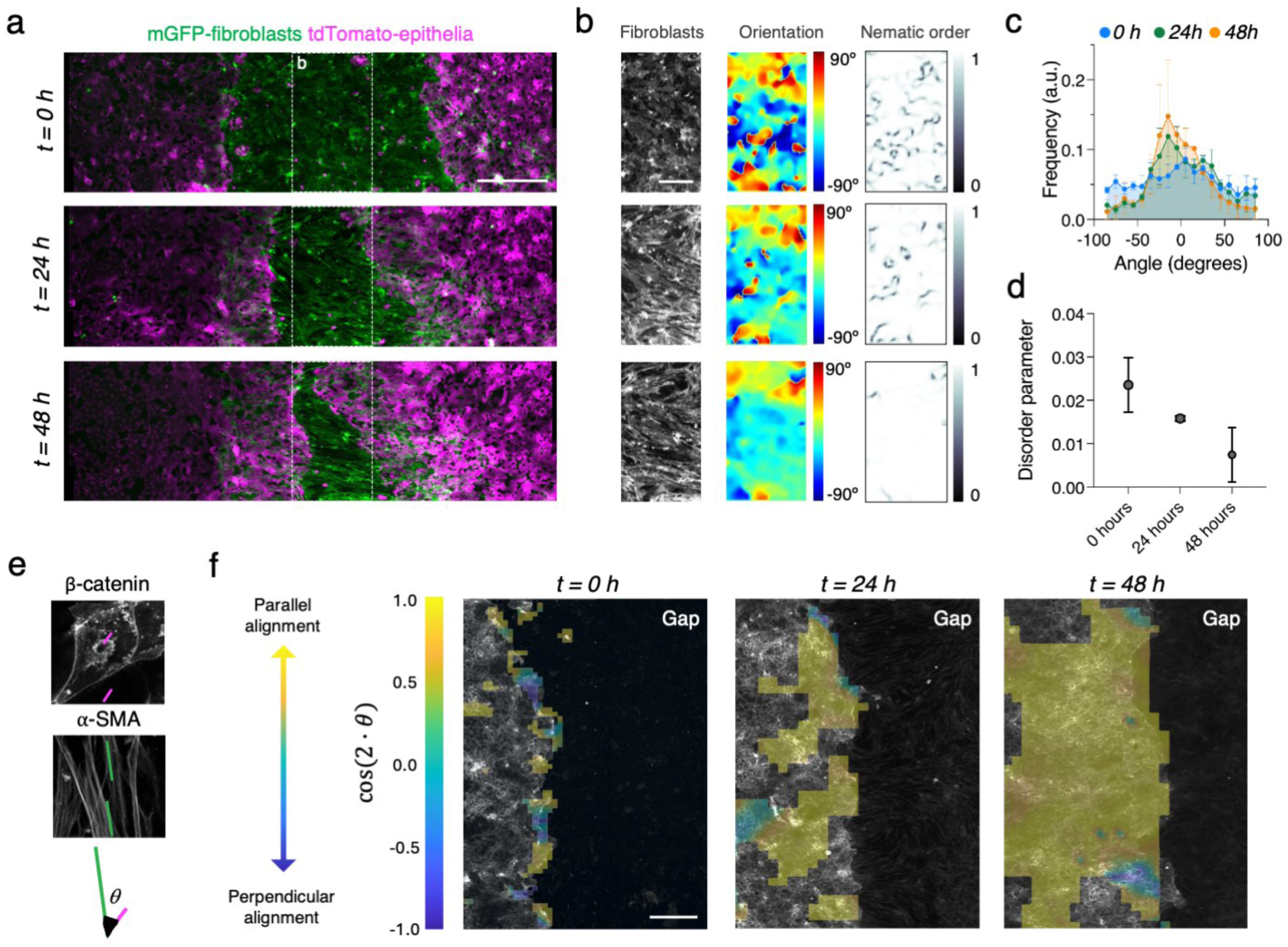
Fibroblasts establish long-range alignment and spatiotemporal co-orientation with epithelial cells during gap closure. (a) Sum intensity projections of timeframes corresponding to 0, 24 and 48h upon barrier removal of the + fibroblasts condition. Scale bars: 500 µm. (b) Representative orientation and nematic order maps of fibroblasts in the central region of the gap (white box in a). Scale bar: 250 µm. (c) Angular distribution of fibroblast orientation at 0, 24 and 48h upon barrier removal. (d) Fibroblasts’ disorder parameter over time. (e) Schematics depicting the dominant directions (w) for each cell type and their angular difference q, from which the correlation index (cos(2·q)) is defined. (f) Correlation maps at different time points (middle and right panels). Yellow stands for regions with parallel alignment between the two cell types and blue for regions with perpendicular alignment. Regions with no color indicate that at least one of the two cell types did not have a defined dominant direction. Scale bar: 500 µm. Immunostained samples from N = 2 independent experiments were used.

At the same time, epithelial cells were becoming oriented in the direction of migration as well. As it can be seen in segmented monolayers (Figure S7), outlines of individual epithelial cells show that cell shape became progressively elongated and oriented in the direction of gap closure. Thus, we next analyzed the temporal co-orientation of cells in the coculture during the gap closure by immunofluorescence using ɑ-SMA and β-catenin antibodies to visualize fibroblasts and IECs, respectively. The dominant directions of fibroblasts and the IECs were computed by dividing the images in subregions (roughly of the area of an IEC) and determining their dominant directions (ω) from the spatial gradient of the fluorescent signal for β-catenin and for α-SMA (Figure 5e). We then calculated the angle (θ) between the dominant directions of fibroblasts and IECs and plotted the value of cos(2·θ) (referred as correlation index, Figure 5f) for each subregion, generating correlation maps indicative of the alignment between the two cell types. The correlation analysis revealed that immediately after removing the barrier (t = 0 h), the alignment between fibroblasts and IECs was weak and restricted to the regions adjacent to the migration front (Figure 5f). However, as time went by (at 24 and 48 h), fibroblasts near the gap were more numerous, elongated (Figure 4a) and aligned with the IECs, becoming predominantly coaligned in the correlation map (Figure 5f, t = 24 and t = 48 h). This alignment extended further into the epithelium monolayer over time. Importantly, the accumulation of intestinal fibroblasts with high levels of α-SMA^+^ stress fibers (Figure 4a) and the coalignment of fibroblasts and IECs matched temporally and spatially with the increased directionality of the epithelial trajectories towards closing the gap. This evidences the active role of fibroblasts, which appear to coordinate with IECs for an efficient epithelial migration.

### Aligned ECM fibers deposited by fibroblasts support directional epithelial migration

In vivo, intestinal fibroblasts remodel the tissue matrix during wound healing through collagen deposition and metalloprotease expression.^9,44–48^ Thus, we investigated whether this matrix deposition could contribute to the enhanced efficiency in epithelial gap closure observed in the physical presence of fibroblasts. First, we evaluated ECM protein deposition in the cocultures. Immunostaining of β-catenin, α-SMA, and collagen IV revealed organized collagen-path depositions accumulating between the fibroblast layer and the epithelium (Figure 6a, b), following the orientation patterns observed for the fibroblasts. To examine this further, we performed immunostaining for fibronectin, laminin, and collagen IV near the migration front in the gaps of conditions with a continuous layer of fibroblasts (+fibroblasts). There, all three ECM components were abundantly present and organized into fiber-like structures (Figure 6c) aligned with the direction of epithelial migration (Figure 6d). So, when fibroblasts are in the gap, they align themselves together with fibronectin, laminin and collagen IV fibers parallel to the direction of epithelial migration. These findings suggest that the long-range organization of fibroblasts along the axis of epithelial migration enables the deposition of aligned ECM fibers, which in turn may guide and reinforce directed epithelial movement during gap closure.

**Figure 6.**
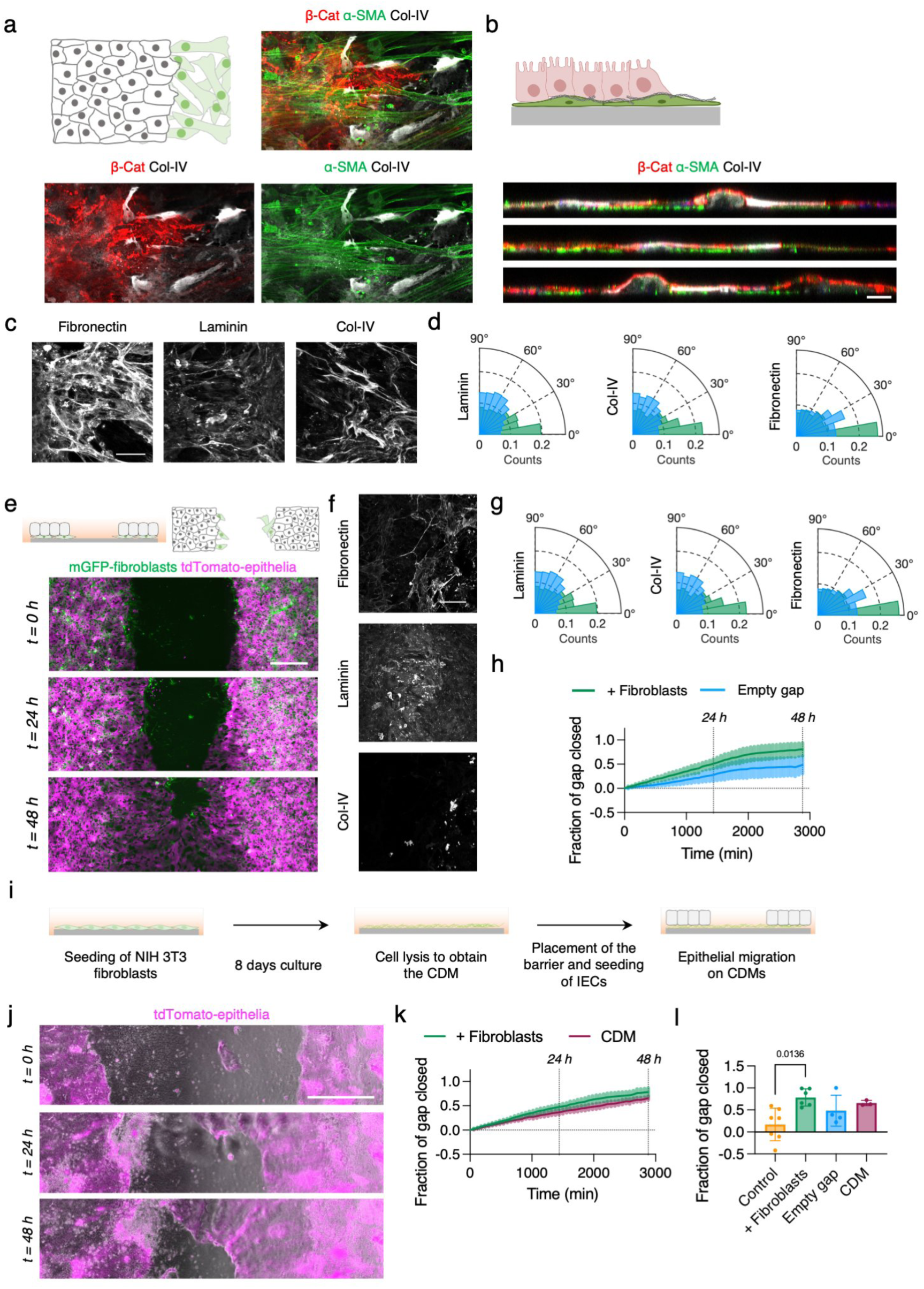
Fibroblasts deposit aligned ECM fibers that support directed epithelial migration during gap closure. (a) Representative fluorescence microscopy images corresponding to immunostaining of β-Catenin (β-Cat), alpha-smooth muscle actin (ɑ-SMA), and collagen-IV (Col-IV) in + fibroblasts condition. Scale bar: 25 µm. (b) Orthogonal cross-sections of the coculture. Scale bar: 10 µm. (c) Sum intensity projections of + fibroblasts condition at 24h upon barrier removal stained for fibronectin, laminin or col-IV. (d) Polar histograms of the fibbers’ orientation of each protein. 0° corresponds to parallel alignment to the migration direction and 90°, perpendicular. (e) Schematics of the experimental set-up of the “empty gap” condition. Sum intensity projections of timeframes corresponding to 0, 24 and 48h upon barrier removal of the empty gap condition. Scale bar: 500 µm. (f) Sum intensity projections of empty condition at 24h upon barrier removal stained for fibronectin, laminin or col-IV. (g) Polar histograms of the fibbers’ orientation of each protein. 0° corresponds to parallel alignment to the migration direction and 90°, perpendicular. (h) Fraction of gap closed over time in + fibroblasts and empty cap conditions. Mean ± SEM. N = 6 and N = 4 independent experiments respectively. (i) Schematics of the preparation of CDMs and the experimental layout for the migration experiments. (j) Snapshots of the live-imaging of mT/mG organoid-derived cells migrating on NIH 3T3 fibroblasts CDMs at 0, 24 and 48h upon barrier removal. Scale bar: 500 µm. (k) Fraction of gap closed along time in + fibroblasts and CDM conditions. Mean ± SEM. N = 6 and N = 3 independent experiments respectively. (l) Fraction of gap closed at 48 in control, + fibroblasts, empty gap and CDM conditions. Mean ± SD. N = 7, N = 6, N = 4 and N = 3 independent experiments for control, + fibroblasts, empty gap and CDM conditions, respectively. Statistical significance was assessed using a Kruskal-Wallis test.

To further test this hypothesis, we designed an additional experimental setup. We compared epithelial cell migration on an intact monolayer of fibroblasts (+ fibroblasts condition) with a scenario where the fibroblast monolayer was interrupted in the gap region (empty gap condition, Figure 6e). Therefore, epithelial cells would be in close contact with the fibroblasts and exposed to their secreted signals, but there should be no long-range order nor secretion of proteins in the gap region far from the epithelial front. To create this setup, we first placed the elastomeric barrier on the substrate and then seeded primary fibroblasts, spatially confining them to both sides of the barrier. Once the fibroblasts had formed a monolayer, we seeded the IECs until they also formed monolayer and we removed the barrier, creating a gap empty of fibroblasts (Figure S8a). Upon removal of the barrier, both cell types started migrating towards the gap simultaneously (Movie 13). We then checked for the amount and distribution of ECM proteins in the case of an empty gap. Signal was dimmer and appeared more globular (Figure 6f). We then checked the alignment of fibers and fibroblasts with the migration front and discovered that, for the empty gap condition, the alignment was lost in ECM proteins and in fibroblasts near the migration front (Figure 6g). This came along with epithelial cells reaching a lower fraction of gap closure when compared to when fibroblasts were present in the gap (+ fibroblasts condition) (Figures 6h). Moreover, net distances were shorter, and the alignment of the trajectories was reduced when the gap was initially devoid of fibroblasts, thus leading to a less persistent and less directed migration (Figure S8b and c). These results suggest that the directional migration of epithelial cells is guided by the presence of aligned protein fibers secreted by the fibroblasts present in the gap region.

Finally, to isolate the contribution of secreted ECM proteins from the paracrine signals produced by fibroblasts that induce a WAE-like phenotype in epithelial cells, we employed fibroblast-derived cell-derived matrices (CDMs) (Figure 6i). Briefly, a confluent monolayer of NIH 3T3 fibroblasts was cultured for 8 days to allow ECM deposition. Cells were then lysed, leaving the secreted ECM stably adhered to the surface.^49,50^ Then we placed the elastomeric barrier and seeded the IECs onto the CDM. Once confluent, the barrier was removed, and the epithelial monolayer was allowed to migrate on the fibroblast-derived ECM.

Epithelial monolayers on CDMs migrated cohesively toward the gap, without exhibiting the hole formation typically observed in control condition (Figure 6j and Movie 14). Interestingly, migrating IECs exerted mechanical forces that induced long-range deformations of the CDM across the gap, even beyond the immediate migration front. When quantifying gap closure over time, IEC monolayers on CDMs showed slightly lower efficiency compared to those migrating on living fibroblasts (Figure 6k), but outperformed the empty gap condition (Figure 6l). These results indicate that the fibroblast-derived ECM alone, when distributed across the gap, is sufficient to enhance epithelial gap closure, even in the absence of paracrine signals or cell contact.

## Discussion

Fibroblasts engage in physical crosstalk with epithelial cells across a wide range of biological contexts, including development,^2,24^ wound healing,^3–9^ and cancer progression.^51–53^ While the paracrine role of intestinal fibroblasts has been extensively characterized, the physical contribution of these cells to intestinal epithelial migration and tissue integrity has remained less well understood. Early in vivo observations suggested that colonic fibroblasts might migrate along crypt walls at rates similar to epithelial cells,^54^ hinting at potential synchronized behavior. More recent work has underscored the necessity of mesenchymal signals for epithelial wound repair.^25–28^ Yet, whether and how physical interactions contribute to epithelial migration and coordination had remained unclear.

Here, we developed an in vitro model of the intestinal mucosa that includes both organoid-derived epithelial cells and primary intestinal subepithelial fibroblasts, allowing detailed study of their interaction. This model recapitulates the main epithelial populations from the crypt and villus compartments, as previously described,^55^ and self-organizes into a compartmentalized monolayer architecture.^32^ The fibroblast compartment includes transcriptionally distinct populations that resemble known in vivo stromal subtypes, including telocytes, trophocytes, pericytes, and PDGFRA^low^ fibroblasts, as identified in scRNA-seq atlases of the intestinal stroma.^14^ Although the spatial organization characteristic of fibroblasts in vivo is not preserved and expression of subtype-specific markers spans multiple clusters, the transcriptional diversity observed in our in vitro system closely mirrors the heterogeneity reported in vivo.

To address functional differences in epithelial behavior across conditions, we used elastomeric barriers to standardize gap formation and systematically compare epithelial responses under distinct stromal contexts: with fibroblast-conditioned medium, on continuous or discontinuous fibroblast monolayers, and on fibroblast-derived ECMs. This approach allowed us to decouple the contributions of paracrine signals, physical contact, and ECM cues in driving epithelial migration.

Using this system, we demonstrate that physical interaction with fibroblasts enhances epithelial directionality, persistence, and integrity during migration. Upon barrier removal, fibroblasts at the gap become activated (α-SMA⁺) and migrate toward the epithelial front. Simultaneously, fibroblasts farther away align along the migration axis and deposit oriented ECM proteins, including fibronectin, laminin, and collagen IV. This large-scale anisotropic matrix organization likely emerges from collective cell behavior: prior work suggests that alignment can arise from cell responses to repeated cell–cell collisions, which reorient cells along a shared axis of movement.^56^ Such long-range ordering extended across hundreds of microns and mirrors nematic alignment phenomena observed in other biological systems, including polar filaments,^57^ the tumor microenvironment,^58^ muscle differentiation,^42,43^ bacterial colonies^59^ and confined monolayers.^60–62^ In our model, aligned fibroblasts were spatially and temporally associated with coaligned epithelial cells, and this coordinated organization correlated with more directed and persistent epithelial migration and more efficient gap closure.

Importantly, this mechanism mimics in vivo repair, where mesenchymal stem cells migrate to wound sites^27^ and epithelial cells differentiate into wound-associated epithelial (WAE) cells. In our model, fibroblast signals induced a WAE-like transcriptional profile, marked by reduced expression of *Ki67* and *Lgr5*, and increased *Cldn4* and *Dpcr1* expression, matching previous WAE cell signatures.^27,28^ Interestingly, this occurred without an increase in overall proliferation, which may instead serve to replace apoptotic cells.^25,63^ We further explored the role of PGE₂–EP4 signaling, known to drive WAE cell identity.^28^ Although adding PGE₂ to control cultures partially induced WAE genes, it failed to fully rescue epithelial migration. Conversely, inhibiting EP4 reversed the WAE-like transcriptional state but did not impair gap closure in co-cultures with fibroblasts. These findings suggest that while PGE₂ promotes a WAE-like transcriptional program, other fibroblast-dependent cues, including ECM contact and mechanical coordination, likely contribute to migration efficiency.

To test the contribution of fibroblast-derived ECM, we used CDMs generated by lysing fibroblasts after matrix deposition. IECs migrating on CDMs performed better than controls, confirming a supportive role for ECM. However, migration remained less efficient than on live fibroblast layers, where dynamic interactions and long-range alignment are present. Moreover, excluding fibroblasts from the gap impaired epithelial migration, even though fibroblasts were present under the epithelium, indicating that fibroblast-fibroblast communication across the gap is required for matrix organization and guidance.

Fibroblasts also play a key role in synchronizing crypt and villus dynamics. A fundamental aspect for effective gap closure is the preservation of epithelial integrity throughout the process.^64^ The advancement of the tissue must occur while maintaining cell-cell contacts. This is an active process involving cell proliferation, delamination of dead cells, changes in cell shape and tissue rearrangements.^65^ In controls and conditioned medium conditions, villus cells migrated while crypt cells remained static, leading to mechanical uncoupling across the tissue. Only in the presence of fibroblasts did crypt cells become migratory, indicating a need for direct stromal contact to coordinate epithelial compartments. This likely explains the better preservation of epithelial integrity in fibroblast-containing cultures. Indeed, epithelial rupture was common in controls, often initiating near large, likely differentiated cells, consistent with previous reports linking hole formation to tensile stress and cell division.^66^ In our model, fibroblasts rapidly localized to rupture sites, where they appeared to assist in re-sealing the monolayer. The prevention and closure of such holes by fibroblasts suggest a previously underappreciated mechanical support role during epithelial regeneration.

In summary, our study has demonstrated the active involvement of primary intestinal fibroblasts in orchestrating epithelial migration *in vitro*. Their presence, accompanied by their self-organization establishing long-range order and the secretion of aligned extracellular matrix fibers, significantly contributes to the enhancement of gap closure and the upholding of epithelial integrity. Beyond their well-known paracrine signaling roles, fibroblasts physically engage with the intestinal epithelium by migrating toward the wound front, aligning across long distances, and depositing oriented ECM fibers that support and direct epithelial closure. These mechanical and spatial cues promote persistent and directional epithelial movement and help preserve tissue cohesion, in part by synchronizing crypt and villus compartment dynamics and facilitating the closure of epithelial gaps. These results deepen our understanding of epithelial-stromal interactions during wound healing and tissue remodeling, and point to fibroblasts not only as sources of biochemical signals, but as spatially organized, mechano-active components of the regenerative niche.

## Materials and Methods

All the resources used in this work are detailed in Table S1.

### Experimental model and study participant details

All experimental protocols involving mice were approved by the Animal care and Use Committee of Barcelona Science Park (CEEA-PCB) and the Catalan government and performed in accordance with their relevant guidelines and regulations. *Lgr5-EGFP-IRES-creERT2* mice have been previously described.^67^ Briefly, *Lgr5-EGFP-IRES-creERT2* mice were generated by homologous recombination in embryonic stem cells targeting the *EGFP-IRES-creERT2* cassette to the ATG codon of the stem cell marker *Lgr5* locus, allowing the visualization of Lgr5^+^ stem cells with a green fluorescent protein (GFP). *Lgr5-EGFP-IRES-creERT2* mice were crossed with the Cre reporter strain Ai9 (RCL-tdT) (JAX-007909) to generate the *Lgr5-EGFP-IRES-creERT2/RCL-tdT* mouse.^41^ Ai9 (RCL-tdT) is designed to have a *loxP*-flanked STOP cassette preventing transcription of the CAG promoter-driven red fluorescent variant (tdTomato), inserted into the *Gt(ROSA)26Sor* locus. Upon Cre-mediated recombination *Lgr5-EGFP-IRES-creERT2/RCL-tdT* mice express robust tdTomato fluorescence in Lgr5-expressing cells and their progeny. *ROSA26-creERT2;mT/mG* (henceforth mT/mG) mouse model^68^ consists of a cell membrane-targeted, two-color fluorescent Cre-reporter allele. Prior to Cre recombination, cell membrane-localized tdTomato (mT) is expressed in all tissues. Upon the activation of Cre recombinase through tamoxifen, all cells (and future cell lineages derived from these cells) express cell membrane localized GFP (mG) instead of mT.

### Method details

#### Intestinal crypts isolation and culture

Intestinal crypts from *Lgr5-EGFP-IRES-creERT2/RCL-tdT* and *ROSA26-creERT2;mT/mG* mice were isolated as previously described.^41,69^ Briefly, small intestines were flushed with PBS and cut longitudinally. Villi were mechanically removed, and intestinal crypts were isolated by incubating the tissue with PBS containing 2 mM EDTA for 30 minutes at 4°C. The digestion content was filtered through a 70 µm pore cell strainer (Biologix Research Co.) to obtain the crypt fraction. Crypts were plated in Matrigel^®^ drops and supplemented with basic medium: advanced DMEM/F12 plus 1% Glutamax, 1% HEPES, Normocin (1:500), 2% B27, 1% N2, 1.25 mM N-acetylcysteine, supplemented with recombinant murine EGF (100 ng mL^−1^), recombinant human R-spondin 1 (200 ng mL^−1^), and recombinant murine Noggin (100 ng mL^−1^), CHIR99021 (3 µM) and valproic acid (1 mM) to formulate the ENRCV medium.^70^ The medium was changed every 2 to 3 days. The first 4 days of culture the Rho kinase inhibitor Y-27632 was added to the culture. Outgrowing crypts were passaged twice a week and organoid stocks were maintained for up to 4 months.

#### Generating Lgr5-EGFP/RCL-tdT intestinal organoids by in vitro tamoxifen dependent creERT2 induction

*Lgr5-EGFP-IRES-creERT2/RCL-tdT* small intestinal organoids were treated *in vitro* with 100 nM of 4-Hydroxitamoxifen (4-HT) for 48 h to induce the expression of tdTomato in Lgr5^+^ cells. Treated intestinal organoids were mechanically and enzymatically digested as described above and cell sorted to obtain pure GFP^+^ and tdTomato^+^ cell populations. Sorted cells were cultured in Matrigel^®^ drops with ENRCV medium plus the Rho Kinase inhibitor Y-27632 to obtain a new *in vitro* line of intestinal organoids named *Lgr5-EGFP/RCL-tdT,* which expresses a robust tdTomato fluorescence in all cells and GFP fluorescence in Lgr5^+^ stem cells. Outgrowing crypts were passaged once a week and organoid stocks were maintained for up to 4 months.

#### Intestinal organoid digestion to single cells

To obtain organoid-derived intestinal epithelial cells (IECs), fully-grown organoids were subjected to a digestion protocol. Briefly, Matrigel^®^ drops containing organoids were disrupted by pipetting with TrypLE Express1X and transferred to a Falcon tube at 4°C, where mechanical disruption was applied using a syringe with a 23 G 1” needle (BD Microlance 3). Disrupted organoids were further digested by incubating them for 5 to 7 minutes at 37°C with vigorous hand-shaking every minute. Successful digestion to single cells was confirmed via inspection under the microscope.

#### Intestinal fibroblasts’ isolation and culture

Intestinal primary fibroblasts were isolated from the same models used to establish organoids’ culture. Lgr5-GFP- and tdTomato-derived stromal cells exhibited a wild-type phenotype (no fluorescence). mTmG-derived stromal cells exhibited cell membrane-localized tdTomato fluorescence (mT). Intestinal primary fibroblasts were isolated from mouse small intestine by adapting a previously published protocol.^71^ Briefly, once the crypts were isolated, the tissue pieces were further digested by first performing 3 incubations of 10 min with 3 mM EDTA in PBS containing at 37°C shaking. Next, after a washing with PBS, the pieces were incubated with collagenase (100 U mL^-1^) in culture medium for 30 minutes at 37°C shaking. Then, the tissue was centrifuged at 1200 rpm for 5 min and the pellet was resuspended in DMEM (1X) with GlutaMAX™ medium supplemented with 10% FBS, 1% Penicillin-Streptomycin and 1% v/v minimum essential medium non-essential amino acids (primary fibroblasts culture medium). The tissue pieces were cultured in flasks kept at 37°C in a humidified incubator under a 5% CO_2_ atmosphere. After seven days in culture, tissue pieces had attached, and fibroblasts started to come out and attach to the flasks. Plates reached confluency after approximately fourteen days. Primary fibroblasts were passaged with a split ratio of 1:2 or 1:3, by first rinsing the cells with warm PBS and incubating them with Trypsin-EDTA for 5 minutes at 37°C. Next, Trypsin-EDTA was neutralized by adding an equal volume of culture medium and cells were centrifuged at 1200 rpm for 5 min. The supernatant was discarded, the pellet was resuspended in culture medium, and the proportional volume was seeded in a new flask containing warm culture medium. Primary fibroblasts were only passaged a maximum of 5 times. mTmG-derived fibroblasts were treated with 4-HT for seven days to convert them from mT+ to mG+ fibroblasts.

#### Preparation of primary fibroblasts conditioned medium

The culture medium used to grow the primary fibroblasts for 4-6 days was collected, centrifuged, and filtered through a 0.22 µm pore size filter (Merck-Millipore). Next, it was supplemented with 2% B27, 1% N2 and 0.25% N-acetylcysteine, obtaining primary fibroblast_CM. To render it suitable for the culture of organoids or organoid-derived cells, the primary fibroblast_CM was supplemented with EGF (100 ng mL^-1^), human R-Spondin 1 (200 ng mL^-1^), Noggin (100 ng mL^-1^), CHIR99021 (3 mM), valproic acid (1 mM), resulting in primary fibroblast_CM/ENRCV.

#### Single cell RNA sequencing of primary fibroblasts

To characterize the different cell populations within the isolated stromal cells, we performed single cell RNA sequencing (scRNAseq). A pool of 3 vials of primary fibroblasts from 3 different isolations at passage 2-4 was prepared to remove the possible variability coming from the mouse selected. Briefly, the 3 vials were centrifuged at 335 rcf during 5 min at 4°C and resuspended in DMEM/F12 with 10% FBS to have a cell density of 300-1000 cells µL^-1^. Cell concentration and viability were determined using a TC20™ Automated Cell Counter (Bio-Rad Laboratories, S.A) upon staining the cells with Trypan blue. Cells were partitioned into Gel Bead-In-Emulsions (GEMs) by using the Chromium Controller system (10X Genomics), with a target recovery of 5000 total cells. cDNA sequencing libraries were prepared using the Next GEM Single Cell 3’ Reagent Kits v3.1, following manufacturer’s instructions. Shortly, after GEM-RT clean up, cDNA was amplified during 12 cycles and cDNA quality control and quantification were performed on an Agilent Bioanalyzer High Sensitivity chip (Agilent Technologies). cDNA libraries were indexed by PCR using the PN-220103 Chromiumi7 Sample Index Plate. Size distribution and concentration of 3’ cDNA libraries were verified on an Agilent Bioanalyzer High Sensitivity chip (Agilent Technologies). Finally, sequencing of cDNA libraries was carried out on an Illumina NovaSeq 6000 using the following sequencing conditions: 28 bp (Read 1) + 8 bp (i7 index) + 0 bp (i5 index) + 89 bp (Read 2), to obtain approximately 20-30.000 reads per cell.

Sequencing reads were processed using CellRanger v.6.1.2.^72^ The output folder was used as input to perform downstream analysis with the R package Seurat 4.0.6 (R.4.1.2). Seurat object was created with the function CreateSeuratObject with the parameter min.cells = 3. To ensure cells of good quality, only cell barcodes within the range of 2000 - 7000 detected genes and < 5 % mitochondrial content were kept for the analysis. Data was normalized with the SCTransform method with default parameters. UMAP was performed with the 30 first principal components, followed by the functions FindNeighbours and FindClusters with default resolution. Cluster markers were identified with the function FindAllMarkers with the options only.pos = TRUE and logfc.threshold = 0.25. Cell type annotation was performed manually looking at the expression of the following genes: *Actg2, Cd52, Pecam1, Lyve1, Rgs5, Pdgfrb, Gfap, Cspg4, Gli1, Myh11, Acta2, Foxl1, Cd34, Pdgfra, Des, Vim, Bmp7, Bmp5, Bmp4, Bmp2, Bmp1, Chrd, Grem1, Wif1, Frzb, Dkk3, Dkk2, Rspo3, Rspo2, Rspo1, Wnt5b, Wnt5a, Wnt4, Wnt2b*.

#### Coculture of intestinal organoids with primary fibroblasts

To test the effect of culturing primary fibroblasts with intestinal organoids, crypts derived from Lgr5-EGFP-IRES-creERT2 intestinal organoids were embedded in Matrigel drops together with primary fibroblasts and cultured with ENRCV medium. Briefly, Matrigel drops containing intestinal organoids were digested with trypsin at 4°C, broken using a syringe with a 23 G 1” needle, and incubated 1 min at RT. Then the pieces of organoids per drop were counted. In parallel, primary fibroblasts were trypsinized, resuspended and counted. Finally, crypt pieces and fibroblasts were mixed in Matrigel at a concentration of 400 crypt pieces and 50.000 fibroblasts in 50 µL of Matrigel to generate the 3D coculture of intestinal organoids and fibroblasts. To test only the paracrine effect of primary fibroblasts, crypts were cultured alone with primary fibroblast_CM/ENRCV medium.

Crypts were also cultured alone using ENRCV medium, ENRCV medium supplemented with 16,16-Dimethyl Prostaglandin E2 1 µM in DMSO (PGE_2_ condition), ENRCV medium supplemented with recombinant human BMP-4 9 nM in HCl, and primary fibroblast_CM/ENRCV medium with EP4 receptor antagonist L-161,982 10 µM in DMSO, ENRCV medium with the corresponding volumes of DMSO and HCl used for PGE_2_, BMP-4, and EP4 receptor antagonist respectively as controls.

Cultures were maintained for 2 days, but occasionally fibroblasts – crypts cocultures were maintained longer, up to 14 days.

#### Quantitative reverse-transcription polymerase chain reaction (RT-qPCR)

Total RNA was extracted from individual Matrigel drops, each cultured in a separate well of a 24-well plate, using 800 µL of TRIzol Reagent Solution® per well. The RNA extraction was achieved following the manufacturer’s instructions of the reagent, and the RNA was resuspended in DEPC treated water. The concentration and purity of the extracted RNA was measured using the Nanodrop 1000 (Thermo Scientific). For RNA-complementary DNA (cDNA) synthesis, 1000 ng of total RNA were first treated with DNAse I Amplification Grade® to remove all genomic DNA. The reverse transcription of the RNA was performed using the iScript™ cDNA first-strand Synthesis kit following the manufacturer’s protocol.

The mRNA transcript levels of the genes of interest were analysed by real-time quantitative PCR using the StepOne Plus real time PCR system (Applied Biosystems). The primers used for each gene (*gapdh*,^73^ *rps18*, *fabp1*,^74^ *ki67*,^75^ *lgr5*,^76^ *cldn4*, *dpcr1*, *cd55b*, *axin2*) are shown in Table S2. The specificity of the primers to recognize correctly the gene sequence, and the absence of non-specific primer bindings were tested in silico. For each sample and gene studied, the analyses were performed in triplicate wells in a 96-well plate. The PCR reactions were carried out in a final volume of 10 µL containing 5 µL of Fast SYBR™ Green Master Mix, 0.5 µL of a mixture of forward and reverse primers (250 nM), 3.5 µL of DEPC water plus 1 µL of diluted cDNA. In addition, three negative controls, a non-template control, a non-reverse transcriptase control and a PCR control (water control), were included and ran in duplicate. The expression level of each gene was calculated with the ΔΔC_T_ method and was analysed relative to the reference genes *gapdh* and *rps18*.

#### Setup of the gap closure models

Ibidi µ-Slides 8 wells (Ibidi GmbH) were coated with Matrigel® to form thin films (< 2 µm) as an extracellular matrix (ECM) surrogate as previously described.^32,35^ Briefly, Ibidi wells were coated with 10 μL cm^-2^ of 3 mg mL^-1^ Matrigel® diluted in DMEM/F-12. To spatially confine cell growth and create a gap where cells could migrate to, we employed PDMS barriers fabricated in-house. For the “control” condition, first the elastomeric barrier was placed and stuck (using Dow Corning® High-Vacuum Grease) on the Matrigel® coated wells. Then, 3.5x10^5^ organoid-derived cells cm^-2^ were seeded with ENRCV medium containing Y-27632 and cells were cultured for 1 day. For the “+ fibr. cond. med.” condition, the elastomeric barrier was placed as in the control condition, then organoid-derived cells were seeded with primary fibroblast_CM/ENRCV medium containing Y-27632, and cells were cultured for 1 day. For the “+ fibroblasts” condition, first, 3x10^4^ primary fibroblasts cm^-2^ were seeded on the Matrigel® coated well and cultured with primary fibroblasts culture medium for 1 day, then the elastomeric barrier was placed, and 3.5x10^5^ organoid-derived cells cm^-2^ were seeded on top with ENRCV medium containing Y-27632 for 1 day more. For the “empty gap” condition, first the elastomeric barrier was placed and stuck on the Matrigel® coated wells. Then, 3x10^4^ primary fibroblasts cm^-2^ were seeded on the Matrigel® coated well and cultured with primary fibroblasts culture medium for 1 day, then 3.5x10^5^ organoid-derived cells cm^-2^ were seeded on top with ENRCV medium containing Y-27632 for 1 day more.

For the CDM condition, Ibidi wells were incubated with 1% Gelatin from porcine skin solution in PBS for 1h at 37°C, afterwards the gelatin solution was removed and gently rinsed with filtered PBS twice, the Gelatin coating was crosslinked with 1% glutaraldehyde solution for 30 min at RT, the glutaraldehyde solution was removed and rinsed twice with filtered PBS. Finally, glutaraldehyde was blocked for 20 min at RT with 1M glycine in PBS, the excess was removed and rinsed twice with filtered PBS. Subsequently, NIH3T3 fibroblasts were seeded at high density (10^6^ cells/cm^2^). The culture medium consisted of DMEM with GlutaMAX, 1% sodium pyruvate, 1% Pen-Strep and 10% FBS was supplemented with 50 μg/mL L-ascorbic acid and changed every day. The culture was maintained for 8 days. Cells were removed by a lysis medium consisting of 20 mM NH_4_OH and 0.5% Triton in PBS after two washing steps with PBS. The pre-warmed lysis medium was carefully pipetted in the well and incubated for up to 10 min at 37°C in the incubator. PBS solution was added and the CDM stored at 4°C. The day after, the PBS solution was carefully changed three times to remove residues of Triton. The matrices were covered with PBS and stored until use at 4°C. Then, the elastomeric barrier was carefully placed and stuck on the CDMs. Then, 3.5x10^5^ organoid-derived cells cm^-2^ were seeded with ENRCV medium containing Y-27632 and cells were cultured for 1 day.

After the indicated times, the migration assay was initiated by carefully removing the elastomeric barrier, washing with warm PBS and adding new corresponding medium without Y-27632. In +PGE_2_ experiments, ENRCV media supplemented with 1 µM PGE2 was added to “control” conditions after the barrier removal. And for the + EP4i experiments, ENRCV media supplemented with 10 µM EP4 receptor inhibitor was added to “+fibroblasts” conditions after the barrier removal.

#### Immunostaining and image acquisition of fixed samples

Cells in Ibidi µ-Slides wells were fixed with 10% neutralized formalin, permeabilized with 0.5% Triton X-100 for 30 minutes, and blocked with 1% BSA, 3% donkey serum, and 0.2% Triton X-100 in PBS for 2 h at RT. Samples were then incubated with the primary antibodies against α-smooth muscle actin, β-catenin, cleaved caspase 3, desmin, vimentin, FOXL1, PDGFRα, Ki-67, fibronectin, laminin, collagen IV, cytokeratin 20, Lysozyme, Wnt3a, EphB2 and Fabp1 (Table S3) overnight at 4°C followed by several PBS washings. Next, samples were incubated with the adequate secondary antibodies (Table S4) plus DAPI (1:1000, Thermo Fisher Scientific) and rhodamine-phalloidin (1:200, Cytoskeleton) for 2 h at RT. Finally, samples were washed with PBS and mounted with Fluoromount G (Southern Biotech). Fluorescence images were acquired using a confocal laser scanning microscope (LSM 800, Zeiss) with a 10x objective (NA = 0.3, WD = 2.0) or 20x objective (NA=0.8, WD=0.55). The laser excitation and emission light spectral collection were optimized for each fluorophore. The pinhole diameter was set to 1 Airy Unit (AU).

#### Live imaging

To optimally track the epithelial cells, single cells derived from tdTomato organoids treated with 4-HT (all cells tdTomato+) and from Lgr5-GFP organoids (only stem cells GFP+) were mixed in a 2:1 ratio, respectively, to reduce the number of fluorescent cells and thus ease cell detection and therefore tracking. Similarly, primary fibroblasts isolated from mTmG mice and treated with 4-HT (mG+) were used in the time-lapse experiments. To register the migration, an Axio Observer 7 epifluorescence inverted microscope (Zeiss) with a 10x objective or a LEICA Thunder (Leica) with 10x objective (NA=0.32, WD= 11.13) or 20x objective (NA=0.4, WD=7.5-6.2), employing temperature (37°C), relative humidity (95%), and CO_2_ (5%) control were used. Phase contrast, 546 and 488 channels were employed. Images were acquired every 10 min up to 48 h of culture.

### Quantification and statistical analysis

#### Fraction of cysts and angle of contact

The fraction of cysts respect to total number of organoids (cysts and budding organoids) was calculated by manual counting of control n = 186 organoids of N = 7 independent experiments, + fibr. cond. med. n = 151 organoids of N = 6 experiments, and + fibroblasts n = 167 organoids of N = 4 experiments. The angle of contact was measured as the angles between the fibroblast’s long axis and the tangent to the cyst. Measurements were obtained from n = 30 fibroblasts from N = 2 experiments.

#### Frequency of Ki-67^+^ epithelial cells

Confocal fluorescence microscopy images of Ki-67 marker were used to quantify the proliferative cells at t=24h. In Imaris, nuclei were first detected with the Imaris Spot detector (diameter set to 9 μm) by manually adjusting the threshold. Next, we applied a F-Actin mean intensity filter to only analyse nuclei within the epithelial monolayer. Then, positive cells for Ki-67 were filtered by applying a threshold of the mean intensity of the marker. Finally, the percentage of Ki-67^+^ cells respect to the total number of cells was computed. N = 3 independent experiments for control, N = 2 for + fibr. cond. med. and N = 2 for + fibroblasts were analyzed.

#### Fraction of gap closed

In Fiji (http://rsb.info.nih.gov/ij, NIH, USA), the cell-free area was measured for frames of cell migration videos every hour and they were normalized to the final cell-free area to obtain the fraction of gap closed. N = 7 independent experiments for control, N = 3 for + fibr. cond. med., N = 6 for + fibroblasts, N = 3 for +PGE2, N = 4 for + fibroblasts + EP4i, N = 4 for empty gap and N = 3 for CDM, were analyzed.

#### Analysis of holes

Holes appearing in the epithelia during gap closure were analyzed by manually outlining the area of each hole in every time point. Hole area was normalized by the maximum area achieved by each hole during its lifetime. n_control_ = 4 holes, n_fib. cond. med._ = 10 holes, and n_+fib_ = 4 holes, from N ≥ 2 independent experiments were analyzed.

#### Cell migration analysis

The centroid trajectories of Lgr5-EGFP/RCL-tdT cells were tracked using the Manual Tracking Plug-in in Fiji. Data analysis was performed using a custom-made code in Matlab (Mathworks, USA). Cell centroid positions during the experiment were defined as **r**_i_ = **r**(iDt), being Dt the time between consecutive images and **r** a vector. The vector difference between the initial (t = t_0_) and the final point (t = t_f_) is defined as the displacement vector **d** and its module || **d** || as the net displacement. The alignment index of the trajectories is defined as cos(2·a), being a the angle between the displacement vector **d** and the gap direction, defined as the direction perpendicular to the epithelial edge. This index equals 1 when the trajectory is parallel to the direction of the gap and -1 when it is perpendicular. For the cell migration analysis, n_control_ = 132 cells, n_fib. cond. med._ = 138 cells, n_+fib_ = 205 cells, and n_empty_ = 118 cells randomly distributed within the epithelia from N ≥ 3 independent experiments were analyzed. For the crypt and villus trackings, n_control_ = 13 cells, n_fib. cond. med._ = 10 cells, n_+fib_ = 10 cells from N ≥ 3 crypts from N ≥ 3 experiments and n_control_ = 17 cells, n_fib. cond. med._ = 18 cells, n_+fib_ = 6 cells from villus regions of N ≥ 3 experiments. PIVlab 2.37^77^ was used to quantify the displacements of IECs and intestinal fibroblasts on phase contrast time-lapse movies. Briefly, an interrogation window of 23 µm with a step of 11.5 µm was used to perform Particle Image Velocimetry (PIV) either by FFT window deformation or by ensemble correlation. Image sequencing was set as time-resolved and the resulting velocities were filtered using a standard deviation filter (8*STD) and a local median filter (threshold = 3). For the PIV analysis, n = 4 migration fronts from N = 2 independent experiments were used.

#### Cell orientation analysis

Orientation fields were obtained using the OrientationJ^78^ plug-in in Fiji. For fibroblasts orientation and nematic order, the ROIs were analyzed using a structure tensor local window of 20 pix and a grid size of 20 pix. Nematic order was measured as 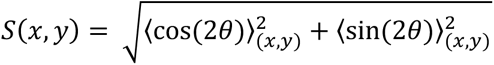, where q is the local orientation of fibroblasts obtained from OrientationJ. Brackets < > denote an average over a local region of interest (x,y) which was defined by 4x4 grid positions each. The fraction of disorder parameter is calculated as the fraction of local regions of interest for which the local nematic order parameter S is below a threshold of 0.5. Images from N = 3 independent experiments were used. For the correlation of cell orientations between fibroblasts and IECs, we proceeded as follows. First, images were pre-processed by subtracting the background (40 pixels rolling ball), then a band-pass filter was applied (limits 3 and 40 pixels), and the background noise was subtracted again (40 pixels rolling ball). The resulting images were processed with OrientationJ using a local window of 40 pixels to obtain a structure tensor on a grid of 40 pixels x 40 pixels. With vector fields and their associated values provided by the plug-in (dominant direction w, energy E and coherence C) we used a custom-made Matlab code to determine the correlation maps. Briefly, we selected the angles with associated energy E > 0.1 and coherency C > 0.05 for each cell type (IEC or intestinal fibroblasts) and discarded the rest. These sets of angles were used to calculate the mean cell orientation and the correlation index. To do so, we collected pairs of dominant angles and obtained the difference between them q = w_IEC_ - w_intestinal fibroblasts_ for each x, y position within the image (note that we only accounted for x, y positions when the dominant angles for both cell types satisfied the conditions for E and C). Then, we defined the correlation index between the two cell types as cos(2·q), which equals 1 when both cells are oriented parallel to each other and -1 when they are perpendicularly oriented. For this analysis, we used immunostained samples from N = 2 independent experiments.

#### Orientation of fibroblasts and deposited proteins

For + fibroblasts and empty gap conditions, orientation fields of fibroblasts at t = 0, 24 and 48 h after barrier removal and of deposited proteins at t=24h were obtained using the OrientationJ plug-in in Fiji using a local window of 2 pixels. Next, the fraction of vector fields respect to the total was computed for each bin spanning 10 degrees. Then, polar histograms were generated in Matlab. For orientation analysis of fibroblasts, we analyzed 14 pictures from N = 5 independent experiments. For orientation analysis of deposited proteins, we analyzed: laminin = 4 ROIs from N = 1, fibronectin = 4 ROIs from N = 2, and collagen IV = 6 ROIs from N = 3.

#### Statistics

No statistical methods were used to predetermine sample size. Measurements were performed on experimental replicates (n) obtained in different independent experiments (N). Data presentation (as Mean value ± standard deviation (SD) or as Mean value ± standard error of the mean (SE)) is defined at the corresponding figure caption. D’Agostino normality test, t-test, and Kruskal-Wallis test were performed using GraphPad Prism 9. Specific values are noted at the corresponding figure captions.

## Data availability

### Lead contact

Requests for further information and resources should be directed to and will be fulfilled by the lead contact: Elena Martínez

### Materials availability

This study did not generate new unique reagents.

### Data and code availability

The scRNA-seq data generated and analyzed during this study have been deposited in https://dataverse.csuc.cat/dataverse/IBEC and are publicly available at https://doi.org/10.34810/data2454. The repository includes raw and processed data files, metadata, and a description of the experimental conditions.

All original code has been deposited on Github (https://github.com/BiomimeticsLab/Comelles_et_al_2025) and is publicly available as of the date of publication.

Any additional information required to reanalyze the data reported in this paper is available from the lead contact upon request.

## Declaration of generative AI and AI-assisted technologies

During the preparation of this work, the author(s) used a large language model (GPT-4o, OpenAI, San Francisco, CA, USA) to improve clarity and grammar. The author(s) reviewed and edited the content as needed and take full responsibility for the content of the publication.

## Acknowledgments

We thank the Martinez Lab members and R.Sunyer for discussions and help, and O. Castillo for assistance in preparing the PDMS barriers.

The authors wish to acknowledge the MicroFabSpace and Microscopy Characterisation Facility, Unit 7 of ICTS “NANBIOSIS” from the CIBER-BBN at IBEC.

Funding for this project was provided by:

European Union Horizon 2020 ERC grant (agreement no. 647863 - COMIET).

Department of Research and Universities of the Generalitat de Catalunya (2021 SGR 01495 to EM and 2021 SGR 001278 to EB).

CERCA Programme/Generalitat de Catalunya.

Networking Biomedical Research Center (CIBER) of Spain. CIBER is an initiative funded by the VI National R&D&i Plan 2008-2011, Iniciativa Ingenio 2010, Consolider Program, CIBER Actions and the Instituto de Salud Carlos III (RD16/0006/0012), with the support of the European Regional Development Fund (ERDF).

Grants PID2023-153116OB-I00 to EB and PID2021-129115OB-I00 to EM funded by MCIN/AEI/10.13039/501100011033 and by “ERDF A way of 2 making Europe”.

Institutional support to CNAG was provided by the Spanish Ministry of Science and Innovation through the Instituto de Salud Carlos III, and by the Generalitat de Catalunya through the Departament de Salut and the Departament de Recerca i Universitats.

The results presented here only reflect the views of the authors; the European Commission is not responsible for any use that may be made of the information it contains.

## Author Contributions

V.F.-M. and E.M. conceived the project. J.C., A.A.-L., V.F.-M., V.A., D.B.-C., A.O.-T. performed experiments. J.C., A.A.-L., and A.O.-T. analyzed data. A.E.-C. performed the analysis of the scRNA-seq data. X.H.-M and E.B. contributed technical expertise, materials and discussion. E.M. acquired the funding for the project. J.C. and A.A.-L. prepared the first draft of the manuscript. All authors discussed the data and assisted in reviewing and editing the manuscript. J.C. and E.M. supervised the project.

