## Supporting Information for "Fibroblast alignment coordinates epithelial migration and maintains intestinal tissue integrity"

**This PDF file includes:**

Supporting text

Figures S1 to S8

Tables S1 to S4

Legends for Movies S1 to S14

SI References

**Other supporting materials for this manuscript include the following:**

Movies S1 to S14

### Supporting Information Text

#### Fibroblast characterization by single-cell RNA sequencing

To characterize the various primary fibroblast populations isolated from mouse intestinal mucosa, we performed single-cell RNA sequencing (scRNA-seq) (Figure S2). A small fraction of the isolated cells, approximately 1%, expressed *Pecam1*, indicating an endothelial identity, with both vascular (*Lyve1*$^{-}$) and lymphatic (*Lyve1*$^{+}$) subtypes represented. Another minor population, around 2%, expressed *Gfap*, a marker typical of neurons and glial cells. Additionally, about 10% of the cells expressed *Cd52*, identifying them as immune cells. The largest fraction, accounting for approximately 85% of the total population, lacked these non-fibroblastic markers but expressed the intermediate filament Vimentin, consistent with a mesenchymal fibroblast identity. Within this fibroblast-enriched compartment, we automatically identified 10 transcriptionally distinct subpopulations.

Although marker expression was not strictly restricted to a single subtype, analysis of canonical and tissue-specific markers revealed distinct clusters corresponding to known stromal cell types. One subset exhibited high expression of *Bmp* genes, such as *Bmp1* and *Bmp4*, along with *Gli1*, a marker not expressed in pericytes,[1] and moderate levels of *Myh11*. This expression pattern is consistent with BMP-producing telocytes or subepithelial myofibroblasts (SEMFs). Within this population, a fraction of cells also expressed *Rgs5* and *Cspg4*, which are typically enriched in pericytes[2] but have also been reported at lower levels in telocytes/SEMFs. Another distinct cluster was enriched for *Dkk3* and *Wnt5a*, with a subset co-expressing *Rspo3*, as previously described for telocytes located at the base of the crypt.[3] These features are consistent with Wnt-modulating telocytes (here referred to as telocytes 1).[4] A distinct subpopulation expressed high levels of *Bmp1* and *Grem1*, with about half of the cells also expressing *Dkk2*; this profile matched that of trophocytes or PDGFRA$^{low}$ stromal cells, known to modulate Wnt and BMP signaling in the crypt niche. Another fibroblast population showed high expression of *Cd34*, *Rspo3*, and *Chrd*, aligning with a well-described PDGFRA$^{low}$ fibroblast subtype (PDGFRA$^{low}$ 1).[5] A separate group expressed high levels of *Pdgfra* and *Wnt4*, consistent with Wnt-secreting telocytes (telocytes 2).

In addition, we identified a population expressing high levels of *Myh11* and *Dkk3*, with moderate *Dkk2* expression, indicative of myocytes with potential Wnt-modulatory capacity. Another subset showed strong *Des* expression but low *Pdgfra* and *Pdgfrb*, suggesting a myocyte/pericyte-like identity. Two additional fibroblast groups displayed gene expression profiles consistent with PDGFRA$^{low}$ fibroblast states (PDGFRA$^{low}$ 2 and 3), although their exact roles remain to be further characterized. Finally, we identified a distinct population with high *Pdgfrb* expression and moderate levels of *Grem1*, consistent with pericyte-like stromal cells that may participate in BMP pathway modulation.

### Figures

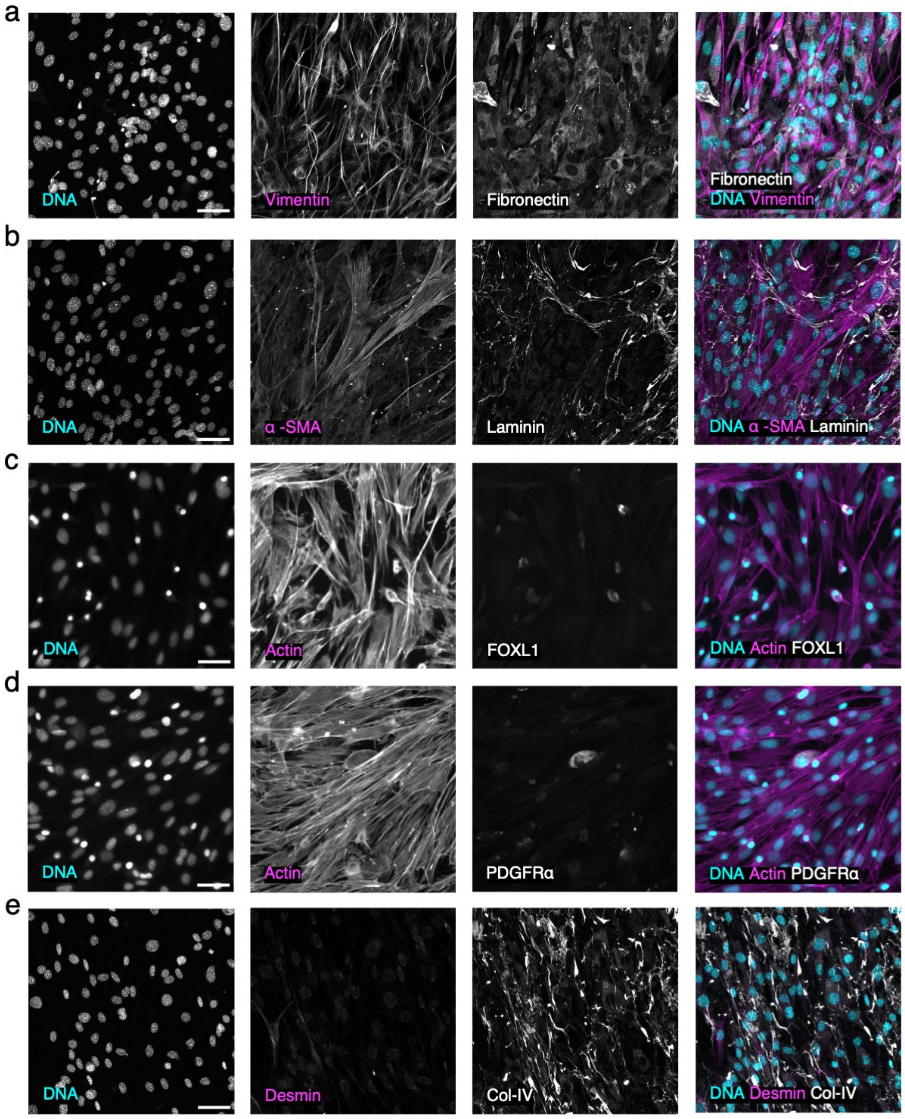


**Figure S1.** Representative fluorescence microscopy images of intestinal fibroblasts corresponding to co-immunofluorescence of (a) DNA, Vimentin and Fibronectin; (b) DNA, a-SMA and laminin; (c) DNA, actin and FOXL1; (d) DNA, actin and PDGFR a; (e) DNA, Desmin and Collagen-IV. Scale bar: 50 μm.

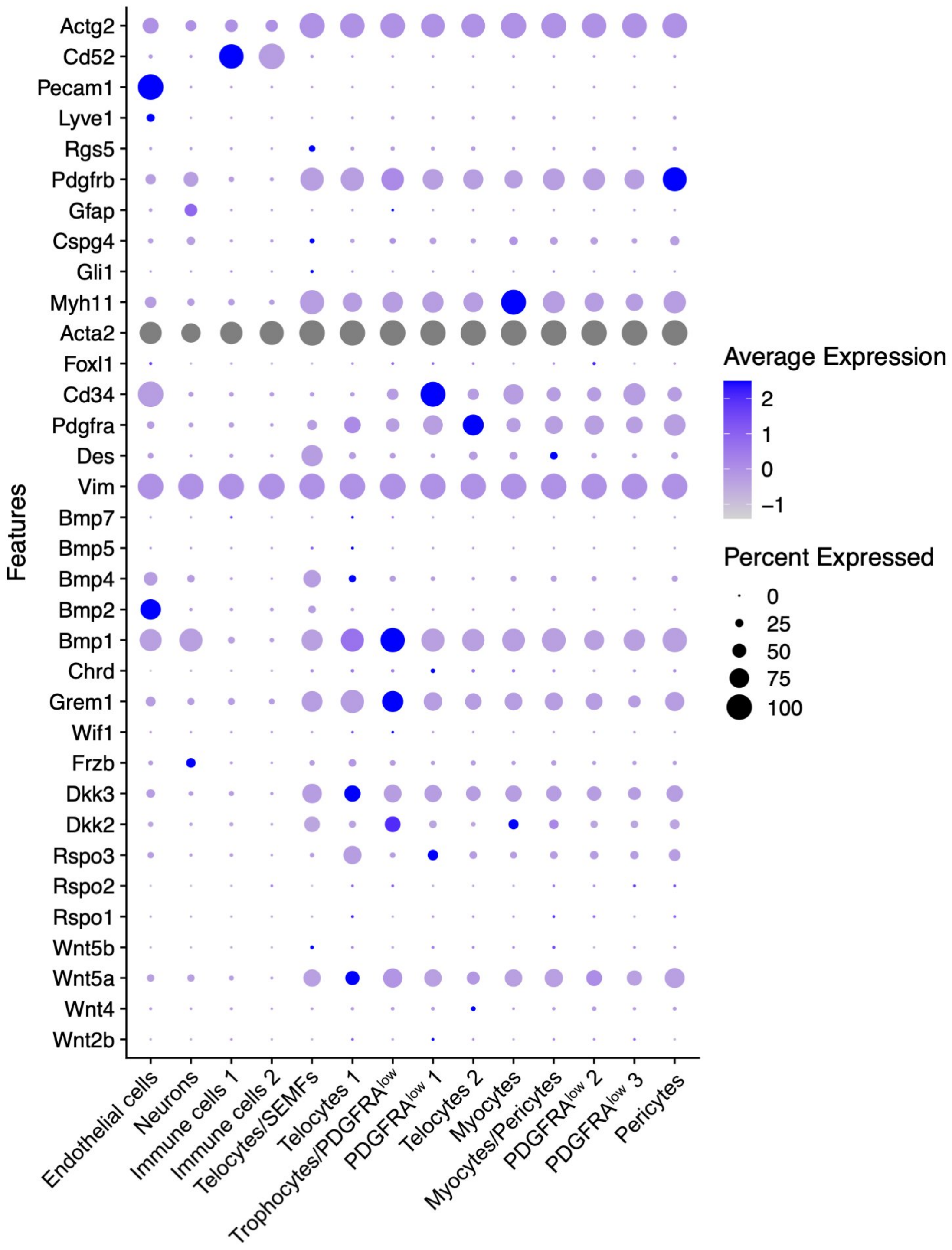


**Figure S2.** Relative expression of known stromal cell markers in cell clusters identified by scRNA-seq. Circle sizes represent the percentage of cells within-cluster expressing that gene and fill colors represent normalized mean expression levels.

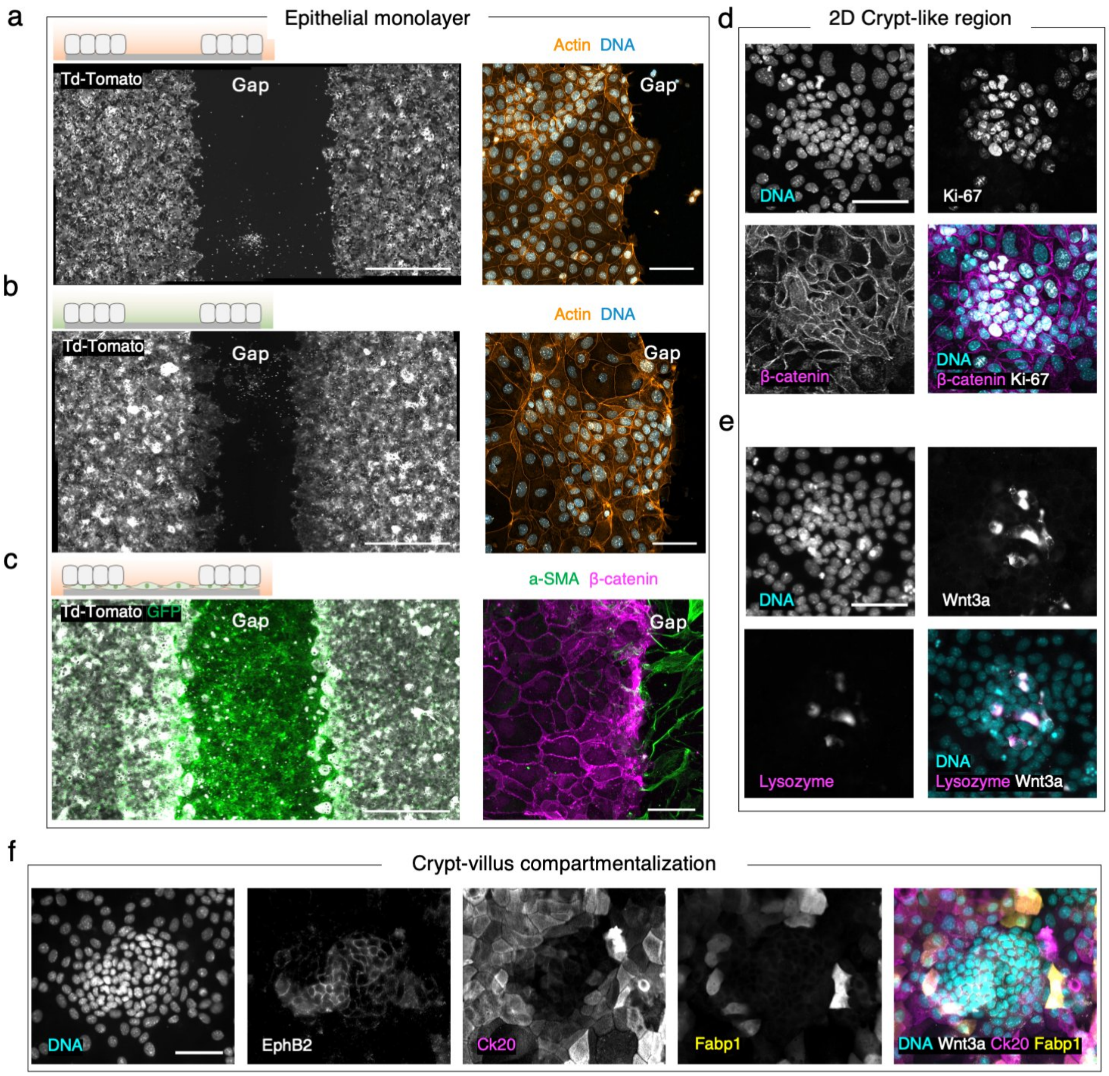


**Figure S3.** Representative confocal images corresponding to co-immunofluorescence of organoid-derived single cell epithelial monolayers. (a) Tile image of the culture well showing the two monolayers (Td-Tomato) separated by the gap and a close view of the gap border (DNA and actin) in control. Scale bars 1 mm and 50 µm. (b) Tile image of the culture well showing the two monolayers (Td-Tomato) separated by the gap and a close view of the gap border (DNA and actin) in +fibr. cond. med. Scale bars 1 mm and 50 µm. (c) Tile image of the culture well showing the two monolayers (Td-Tomato) separated by the gap, where the bottom is covered by a layer of primary fibroblasts (GFP); and a close view of the gap border (a-SMA and β-Catenin) in +fibroblasts. Scale bars 1 mm and 25 µm. (d) Co-immunofluorescence of organoid-derived single cell epithelial monolayers for DNA, Ki-67 and β-Catenin. Scale bar 50 µm. (e) Co-immunofluorescence of organoid-derived single cell epithelial monolayers for DNA, Wnt3a and Lysozyme. Scale bar 50 µm. (f) Co-immunofluorescence of organoid-derived single cell epithelial monolayers for DNA, EphB2, Ck20 and Fabp1. Scale bar 50 µm.

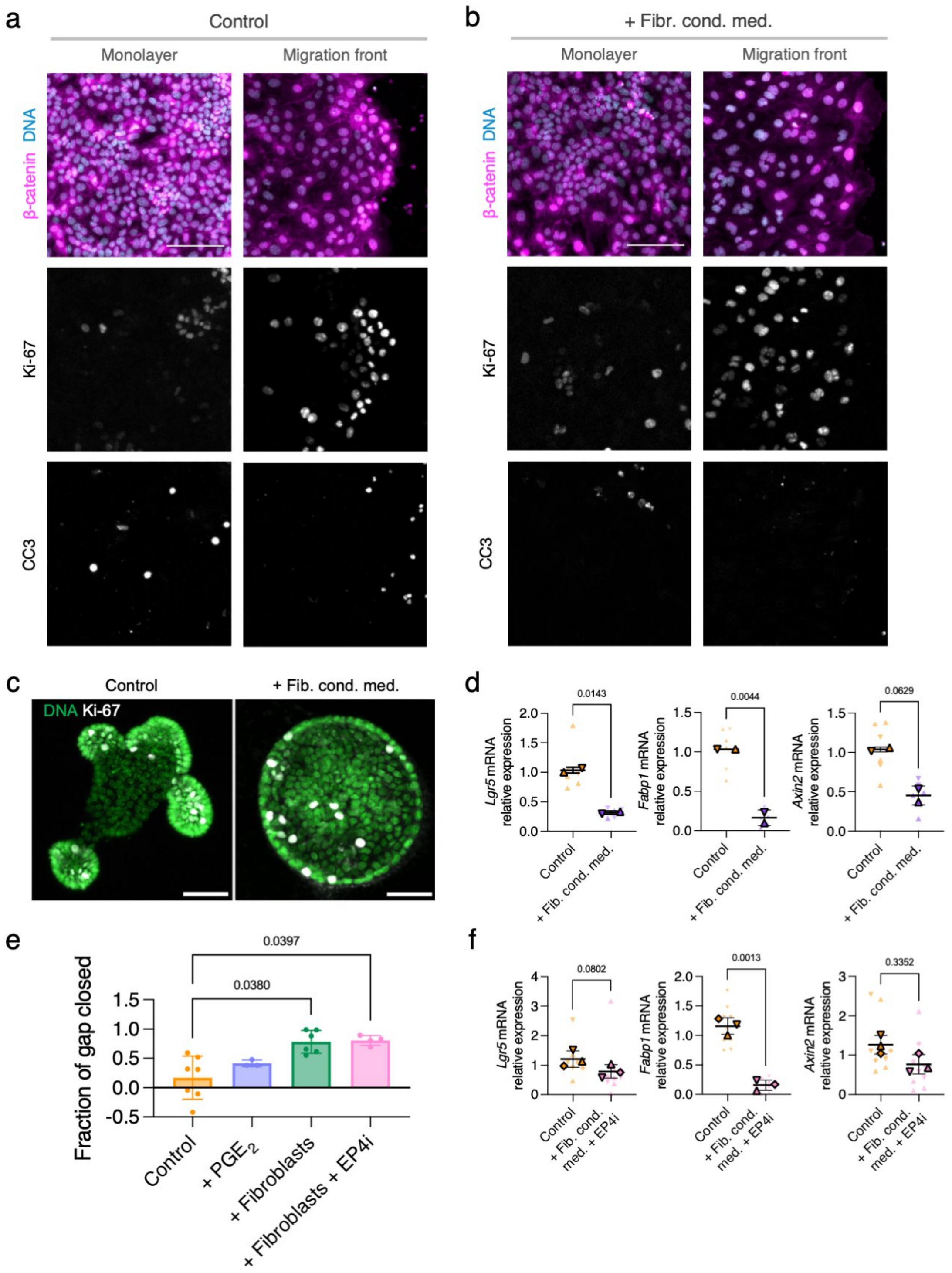


**Figure S4.** (a) Representative sum intensity projections images of epithelial cells in the monolayer region and in the migration front region at 24h upon barrier removal immunostained for DAPI (DNA), β-Catenin, Ki-67 and CC3 in control. (b) Representative sum intensity projections images of epithelial cells in the monolayer region and in the migration front region at 24h upon barrier removal immunostained for DAPI (DNA), β-Catenin, Ki-67 and CC3 in + fibr. cond. med. Scale bar 100 µm. (c) Representative confocal images of crypts grown in 3D Matrigel® drops in control and in + Fibr. cond. med. Scale bars: 50 µm. (d) *Lgr5*, *Fabp1* and *Axin2* mRNA relative expression for control and + fibr. cond. Med 3D Matrigel® drops. Mean ± SD. N = 6 drops from 2 independents experiments per condition. Statistical significance was assessed using a Kruskal-Wallis test. (e) Fraction of gap closed at 48 in control, + PGE2, + fibroblasts and + fibroblasts+EP4i conditions. Mean ± SD. N = 7, N = 3, N = 6 and N = 4 independent experiments respectively. Statistical significance was assessed using a Kruskal-Wallis test. (f) *Lgr5*, *Fabp1* and *Axin2* mRNA relative expression for control and + fibr. cond. med. + EP4i 3D Matrigel® drops. Mean ± SD. N = 9 drops from 3 independents experiments per condition. Statistical significance was assessed using a Kruskal-Wallis test.

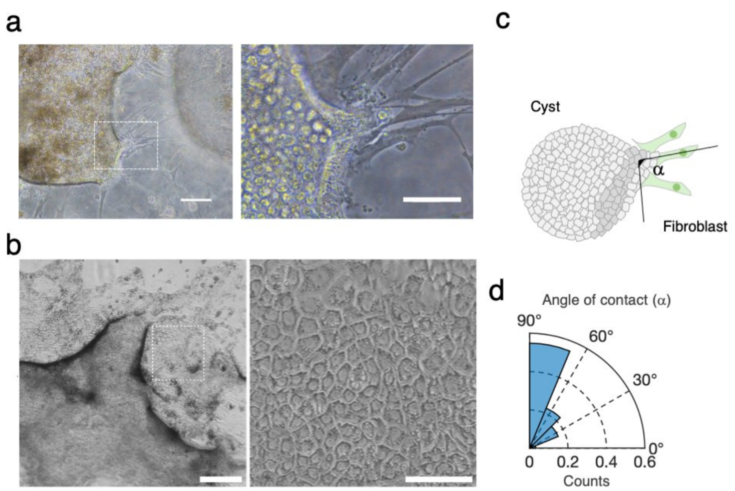


**Figure S5.** (a) Phase contrast image of the expansion of epithelial cysts mediated by fibroblasts. Scale bar = 100 µm. Zoom-in scale bar = 50 µm. (b) Representative bright field microscopy images of + fibroblasts condition after 14 days in culture. Scale bar: 250 µm. Zoom-in. Scale bar: 50 µm. (c) Schematics of the angle of contact between cysts and fibroblasts. (d) Distribution of the angle of contact between cysts and fibroblasts.

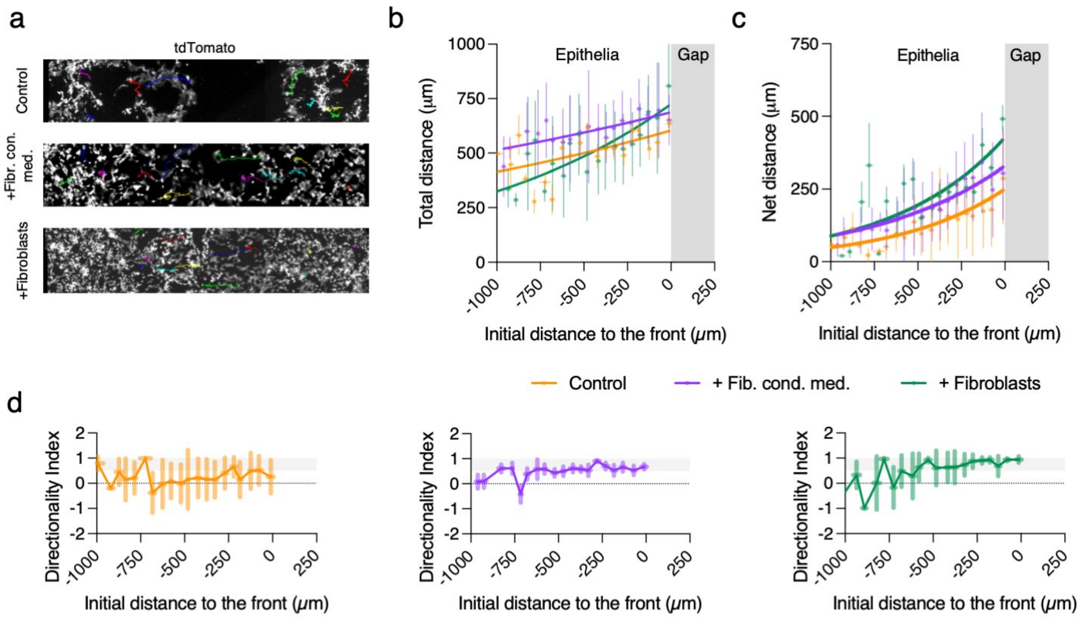


**Figure S6.** (a) Individual cell trajectories for each of the culture conditions (tdTomato organoid-derived cells). (b) Total distance as a function of the initial distance to the migration front for each culture condition. Dots represent the mean value ± SEM, the solid line corresponds to an exponential fitting. The gray region corresponds to the gap. (c) Net displacement as a function of the initial distance to the migration front for each culture condition. Dots represent the mean value ± SEM, the solid line corresponds to an exponential fitting. The gray region corresponds to the gap. (d) Directionality index (cos(2·a), being a the angle between the net displacement vector and the direction perpendicular to the migration front at t = 0) as a function of the initial distance to the migration front for each culture condition. Dots represent the mean value, and the bars represent the SD, the solid line corresponds to a spline-fitting. Gray region corresponds to values that are considered aligned.

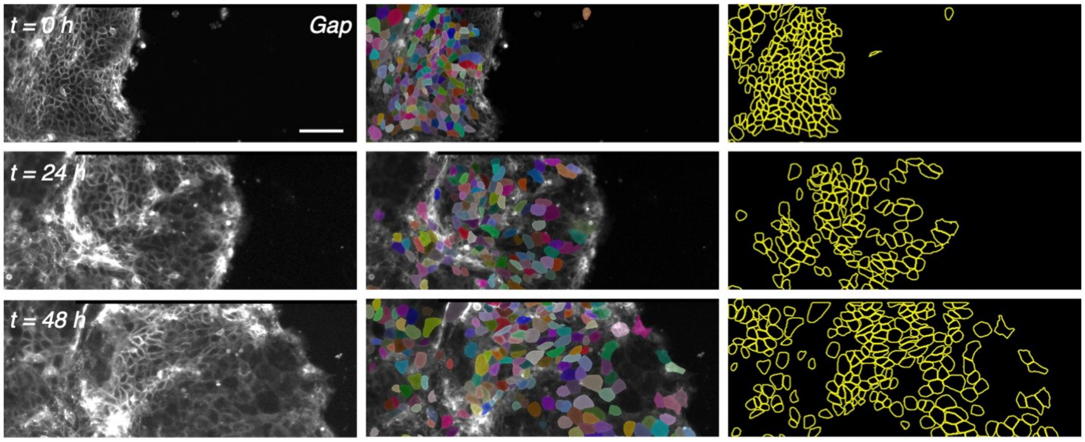


**Figure S7.** Segmentation of mT/mG organoid-derived epithelial cells. From left to right, tdTomato signal of the epithelial monolayer, overlay of the tdTomato signal with the masks for individual cells, and outlines of the segmented cells. Scale bar 100 μm.

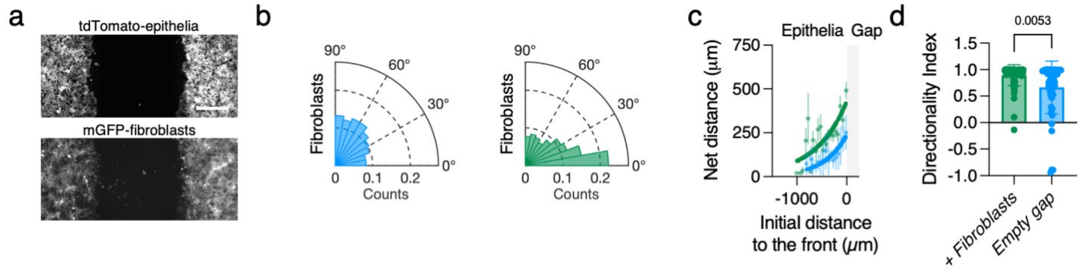


**Figure S8.** (a) Experimental set-up of the "empty gap" condition. Scale bar: 500 μm. (b) Polar histograms of the fibroblasts orientation for empty gap and + fibroblasts conditions. 0º corresponds to parallel alignment to the migration direction and 90º, perpendicular. (c) Net displacement as a function of the initial distance to the migration front. Dots represent the mean value, and bars the SEM, the solid line corresponds to an exponential fitting. Gray region corresponds to the gap outside the epithelium. (d) Directionality index for cells' trajectories within the first 250 μm behind the migration front. Mean + SD and individual values.

**Tables**

**Table S1.** Key resources.

| REAGENT or RESOURCE | SOURCE | IDENTIFIER |
|---|---|---|
| **Antibodies** | | |
| α-smooth muscle actin mouse monoclonal antibody | Abcam | Cat# ab7817<br>RRID: AB_262054 |
| β-catenin goat polyclonal antibody | Biotechne | Cat# AF 1329<br>RRID: AB_354736 |
| β-catenin rabbit polyclonal antibody | Abcam | Cat# ab2365<br>RRID: AB_303014 |
| Cleaved caspase 3 (CC3) rabbit polyclonal antibody | Cell Signaling | Cat# #9661<br>RRID: AB_2341188 |
| Collagen type IV goat polyclonal antibody | Biorad | Cat# 134001<br>RRID: AB_2082646 |
| Desmin rabbit polyclonal antibody | Abcam | Cat# ab15200<br>RRID: AB_301744 |
| Fibronectin rabbit polyclonal antibody | Sigma | Cat# F3648<br>RRID: AB_476976 |
| Ki-67 mouse monoclonal antibody | BD Pharmigen | Cat# 550609<br>RRID: AB_393778 |
| Ki-67 rabbit monoclonal antibody | Abcam | Cat# ab16667<br>RRID: AB_302459 |
| Laminin rabbit polyclonal antibody | Abcam | Cat# ab11575<br>RRID: AB_298179 |
| Vimentin rabbit monoclonal antibody | Abcam | Cat# ab92547<br>RRID: AB_10562134 |

| | | |
|---|---|---|
| FOXL1 mouse monoclonal antibody | Santa Cruz | Cat# sc-130373<br>RRID: AB_2106183 |
| PDGFRα mouse monoclonal antibody | Santa Cruz | Cat# sc-398206<br>RRID: AB_2938643 |
| Cytokeratin 20 mouse monoclonal antibody | Agilent | Cat# M7019<br>RRID: AB_2133718 |
| Lysozyme rabbit polyclonal antibody | Agilent | Cat# A0099<br>RRID: AB_2341230 |
| Wnt3a rabbit recombinant antibody | Abcam | Cat# ab219412<br>RRID: AB_2924383 |
| Wnt3a mouse polyclonal antibody | Abcam | Cat# ab169175<br>RRID: |
| Ephb2 mouse monoclonal antibody | Santa Cruz | Cat# sc-130068<br>RRID: AB_2099958 |
| Ephb2 goat polyclonal antibody | R&D systems | Cat# AF467<br>RRID: AB_355375 |
| Fabp1 rabbit polyclonal antibody | Atlas Antibodies | Cat# HPA028275<br>RRID: AB_10600909 |
| **Chemicals, peptides, and recombinant proteins** | | |
| Ethylenediaminetetraacetic acid | Sigma | Cat# 03690-100ML |
| Matrigel® | Corning | Cat# 354234 |
| Advanced DMEM/F12 | Invitrogen | Cat# 12634-010 |
| Glutamax Supplement | Gibco | Cat# 35050-038 |

| | | |
|---|---|---|
| HEPES solution | Sigma | Cat# H3537 |
| Normocin | Ibian Technologies | Cat# ant-nr-2 |
| B-27 Supplement (50x), minuts vitamin A | Gibco | Cat# 12587-010 |
| N2 supplement | Gibco | Cat# 17502-048 |
| N-Acetyl-L-cysteine | Sigma | Cat# A9165-5G |
| EGF Recombinat Mouse Protein | Gibco | Cat# PMG8041 |
| Recombinant Human R-Spondin 1 protein | R&D Biosystems | Cat# 4645-rs |
| Mouse Noggin Recombinant Protein | Peprotech | Cat# 250-38-250UG |
| CHIR99021 | Stemgent | Cat# 04-0004 |
| Valproic acid sodium salt 98% | Sigma | Cat# P4543-10G |
| Y-27632 | Sigma | Cat# Y0503-1MG |
| 4-Hydroxytamoxifen | Sigma | Cat# H6278 |
| TrypLE Express with Phenol Red | Invitrogen | Cat# 12605 |
| Collagenase from Clostridium histolyticum | Sigma | Cat# C0130-100MG |
| DMEM, High Glucose, GlutaMAX™, Pyruvate | Gibco | Cat# 31966-021 |
| Fetal Bovine Serum (FBS) | Gibco | Cat# A5256701 |
| Penicillin-Streptomycin | Sigma | Cat# P0781-100ML |
| Minimum essential medium non-essential amino acids | Invitrogen | Cat# 11140035 |
| Trypsin 0.05% EDTA | Gibco | Cat# 25300-062 |
| Trypan Blue stain 0,4% | Invitrogen | Cat# T10282 |
| 16,16-Dimethyl Prostaglandin E2 | Biotechne | Cat# 39746-25-3 |
| Dimethyl sulfoxide, >99.7%, Sterile | Sigma | Cat# D2650 |
| BMP-4 | Biotechne | Cat# 314-BP |
| Hydrochloric Acid 37% | Panreac | Cat# 471020 |
| EP4 receptor antagonist L-161,982 | Biotechne | Cat# 2514 |

| | | |
|---|---|---|
| TRIzol™ Reagent Solution® | Thermofisher | Cat# 15596018 |
| DNAse I Amplification Grade® | Sigma | Cat# 10104159001 |
| Fast SYBR™ Green Master Mix | Thermofisher | Cat# 4385610 |
| Dow Corning® High-Vacuum Grease | Corning | Cat# 44224 |
| L-ascorbic acid | Sigma | Cat# 95209 |
| Gelatin from porcine skin | Sigma | Cat# G2500 |
| Glutaraldehyde solution | Sigma | Cat# G6257 |
| Glycine | Sigma | Cat# G8898 |
| $NH_4OH$ | Sigma | Cat# 338818 |
| Triton™ X-100 | Sigma | Cat# X100 |
| Albumin from Bovine Serum (BSA) | Invitrogen | Cat# A23017 |
| Normal Donkey Serum | Abcam | ab7475 |
| DAPI | Invitrogen | Cat# D1306 |
| Rhodamine-phalloidin | Tebu-bio | Cat# PHDR1 |
| Fluoromount-G | Southern Biotech | Cat# 0100-01 |
| Alexa Fluor™ 647 donkey anti-mouse IgG (H+L) | Invitrogen | *Cat#* A-31571 |
| Alexa Fluor™ 568 donkey anti-mouse IgG (H+L) | Invitrogen | *Cat#* A-10037 |
| Alexa Fluor™ 488 donkey anti-mouse IgG (H+L) | Invitrogen | *Cat#* A-21202 |
| Alexa Fluor™ 647 donkey anti-rabbit IgG (H+L) | Invitrogen | *Cat#* A-31573 |
| Alexa Fluor™ 568 donkey anti-rabbit IgG (H+L) | Invitrogen | *Cat#* A-10042 |
| Alexa Fluor™ 488 donkey anti-rabbit IgG (H+L) | Invitrogen | *Cat#* A-21206 |
| Alexa Fluor™ 647 donkey anti-goat IgG (H+L) | Invitrogen | *Cat#* A-21447 |
| Alexa Fluor™ 568 donkey anti-goat IgG (H+L) | Invitrogen | *Cat#* A-11057 |
| Alexa Fluor™ 488 donkey anti-goat IgG (H+L) | Invitrogen | *Cat#* A-11055 |
| **Critical commercial assays** | | |
| Next GEM Single Cell 3' Reagent Kits v3.1 | 10X Genomics | Cat# PN-1000268 |
| iScript™ cDNA first-strand Synthesis kit | BioRad | Cat# 1708890 |

| | | |
|---|---|---|
| **Deposited data** | | |
| Stromal RNA datasets. Raw data | This paper | N/A |
| **Experimental models: Cell lines, organoids and primary cells** | | |
| NIH/3T3 | ATCC | Cat# CRL-1658 |
| *Lgr5-EGFP-IRES-creERT2* mouse intestinal organoids | Barker et al., 2007 | JAX Stock n°: 008875 |
| *EGFP-IRES-creERT2/RCL-tdT* mouse intestinal organoids | Barriga et al., 2017 | N/A |
| *ROSA26-creERT2;mT/mG* mouse intestinal organoids | Muzumdar et al., 2007 | JAX Stock n°: 007576 |
| *Lgr5-EGFP-IRES-creERT2* mouse intestinal fibroblasts | Barker et al., 2007 | JAX Stock n°: 008875 |
| *EGFP-IRES-creERT2/RCL-tdT* mouse intestinal fibroblast | Barriga et al., 2017 | N/A |
| *ROSA26-creERT2;mT/mG* mouse intestinal fibroblasts | Muzumdar et al., 2007 | JAX Stock n°: 007576 |
| **Oligonucleotides** | | |
| *gapdh* F: CTCCCACTCTTCCACCTTCG<br>*gapdh* R: GCCTCTCTTGCTCAGTGTCC | OriGene | Ruiz-Villalba et al., 2017 |
| *rps18* F: CGGAAAATAGCCTTCGCCATCAC<br>*rps18* R: ATCACTCGCTCCACCTCATCCT | OriGene | MP212910 |
| *fabp1* F: GTCAGAAATCGTGCATGAAGGG<br>*fabp1* R: GAACTCATTGCGGACCACTTT | OriGene | Fu et al., 2023 |
| *ki67* F: ATCATTGACCGCTCCTTTAGGT<br>*ki67* R: GCTCGCCTTGATGGTTCCT | OriGene | Santos et al., 2019 |
| *lgr5* F: CCTACTCGAAGACTTACCCAGT<br>*lgr5* R: GCATTGGGGTGAATGATAGCA | OriGene | Xiong et al., 2022 |
| *cldn4* F: CGAGCCCTTATGGTCATCAGCA<br>*cldn4* R: ATGCTTGCCACGATGAACACGG | OriGene | MP202589 |

| | | |
|---|---|---|
| *dpcr1* F: CCTGCTGGTTTTCATTGGCCTG<br>*dpcr1* R: CTGCTCCATCAGGTAAACTGGG | OriGene | MP203665 |
| *cd55b* F: ACGCCTCAGAAACCCACAACAG<br>*cd55b* R: CGTGGTATCAGGTTCATGCTGG | OriGene | MP203452 |
| *axin2* F: ATGGAGTCCCTCCTTACCGCAT<br>*axin2* R: GTTCCACAGGCGTCATCTCCTT | OriGene | MP201191 |
| **Software and algorithms** | | |
| CellRanger v.6.1.2. | 10X Genomics | N/A |
| Seurat 4.0.6 (R.4.1.2) | Butler et al., 2018 | https://github.com/satijalab/seurat |
| RT-qPCR: Step One Plus software (version 2.3) | Applied Biosystems | N/A |
| Zeiss Zen software (version 3.7) | Zeiss | N/A |
| MATLAB version 9.14.02489007 (R2023a) | The MathWorks | N/A |
| PIVlab 2.37 | Thielicke et al., 2014 | https://www.pivlab.de |
| Custom made code | This paper | *https://github.com/BiomimeticsLab/Comelles_et_al_2025* |
| ImageJ Fiji | Schindelin et al., 2012 | https://imagej.net/Fiji/ |
| OrientationJ | Püspöki et al., 2016 | https://bigwww.epfl.ch/demo/orientation/ |
| Leica Application Suite (LASX) | Leica | N/A |
| Imaris software (version 9.1.0) | Bitplane | N/A |
| GraphPad Prism 10 | Dotmatics | N/A |
| **Other** | | |
| Large Language Model (GPT-4o) | OpenAI | N/A |

**Table S2.** Primers used in the qPCR analyses. F: Forward; R: Reverse; Ta: annealing temperature.

| Gene | Primer Sequences (5'-3') | Ta(ºC) | Accession number |
|---|---|---|---|
| ***gapdh*** | F: CTCCCACTCTTCCACCTTCG<br>R: GCCTCTCTTGCTCAGTGTCC | 60 | NM_001289726.2 |
| ***rps18*** | F: CGGAAAATAGCCTTCGCCATCAC<br>R: ATCACTCGCTCCACCTCATCCT | 60 | NM_011296.3 |
| ***fabp1*** | F: GTCAGAAATCGTGCATGAAGGG<br>R: GAACTCATTGCGGACCACTTT | 60 | NM_017399.5 |
| ***ki67*** | F: ATCATTGACCGCTCCTTTAGGT<br>R: GCTCGCCTTGATGGTTCCT | 60 | NM_001081117.2 |
| ***lgr5*** | F: CCTACTCGAAGACTTACCCAGT<br>R: GCATTGGGGTGAATGATAGCA | 60 | NM_010195.2 |
| ***cldn4*** | F: CGAGCCCTTATGGTCATCAGCA<br>R: ATGCTTGCCACGATGAACACGG | 60 | NM_009903.2 |
| ***dpcr1*** | F: CCTGCTGGTTTTCATTGGCCTG<br>R: CTGCTCCATCAGGTAAACTGGG | 60 | NM_001033366.3 |
| ***cd55b*** | F: ACGCCTCAGAAACCCACAACAG<br>R: CGTGGTATCAGGTTCATGCTGG | 60 | NM_007827.3 |
| ***axin2*** | F: ATGGAGTCCCTCCTTACCGCAT<br>R: GTTCCACAGGCGTCATCTCCTT | 60 | NM_015732.4 |

**Table S3.** List of primary antibodies used for immunostainings.

| NAME | TYPE | SOURCE | DILUTION |
|---|---|---|---|
| Anti-α-smooth muscle actin | Mouse Monoclonal IgG2a | Abcam ab7817 | 1:200 |
| Anti-β-catenin | Polyclonal Goat IgG | AF1329 Biotechne | 1:200 |
| Anti-β-catenin | Rabbit IgG pAb | Abcam ab2365 | 1:100 |
| Anti-cleaved caspase 3 (CC3) | Rabbit pAb | Cell Signaling #9661 | 1:200 |
| Anti-collagen type IV | Goat | Biorad 134001 | 1:200 |
| Anti-desmin | Rabbit pAb | Abcam ab15200 | 1:200 |
| Anti-fibronectin | Rabbit | Sigma F3648 | 1:200 |
| Anti-Ki67 | Mouse IgG1 mAb | BD Pharmingen 550609 | 1:100 |
| Anti-Ki67 | Rabbit IgG mAb | Abcam ab16667 | 1:100 |
| Anti-laminin | Rabbit Polyclonal | Abcam ab11575 | 1:200 |
| Anti-vimentin | Rabbit mAb | Abcam ab92547 | 1:200 |
| Anti-FOXL1 | Mouse IgG monoclonal | Santa Cruz sc-130373 | 1:100 |
| Anti-PDGFRα | Mouse IgG monoclonal | Santa Cruz sc-398206 | 1:50 |
| Anti-cytokeratin 20 | Mouse monoclonal (Ks20.8) | Dako M701901 | 1:100 |
| Anti-Lysozyme | Rabbit Polyclonal | Dako A0099 | 1:100 |
| Anti-Wnt3a | Rabbit Recombinant (EPR21889) | Abcam ab219412 | 1:200 |
| Anti-Wnt3a | Mouse Polyclonal | Abcam ab169175 | 1:200 |
| Anti-EphB2 | Mouse monoclonal | Santa Cruz Biotech Sc-130068 | 1:200 |
| Anti-EphB2 | Polyclonal Goat IgG | R&D systems 467B2 | 1:100 |
| Anti-FABP1 | Rabbit Polyclonal IgG | Atlas Antibodies HPA02875 | 1:50 |

**Table S4.** List of secondary antibodies used for immunostainings.

| TYPE | MADE IN | NAME | SOURCE | DILUTION |
|---|---|---|---|---|
| Anti-mouse | Donkey | Alexa Fluor™ 647 donkey anti-mouse IgG (H+L) | Invitrogen A-31571 | 1:500 |
| | Donkey | Alexa Fluor™ 568 donkey anti-mouse IgG (H+L) | Invitrogen A-10037 | 1:500 |
| | Donkey | Alexa Fluor™ 488 donkey anti-mouse IgG (H+L) | Invitrogen A-21202 | 1:500 |
| Anti-rabbit | Donkey | Alexa Fluor™ 647 donkey anti-rabbit IgG (H+L) | Invitrogen A-31573 | 1:500 |
| | Donkey | Alexa Fluor™ 568 donkey anti-rabbit IgG (H+L) | Invitrogen A-10042 | 1:500 |
| | Donkey | Alexa Fluor™ 488 donkey anti-rabbit IgG (H+L) | Invitrogen A-21206 | 1:500 |
| Anti-goat | Donkey | Alexa Fluor™ 647 donkey anti-goat IgG (H+L) | Invitrogen A-21447 | 1:500 |
| | Donkey | Alexa Fluor™ 568 donkey anti-goat IgG (H+L) | Invitrogen A-11057 | 1:500 |
| | Donkey | Alexa Fluor™ 488 donkey anti-goat IgG (H+L) | Invitrogen A-11055 | 1:500 |

**Movie S1 (separate file).** Gap closure experiment in 'control' condition. tdTomato signal from the *Lgr5-EGFP/RCL-tdT* organoid-derived epithelial cell monolayers. Time hh:mm. Scale bar: 500 µm.

**Movie S2 (separate file).** Gap closure experiment in + fibroblast condition medium. tdTomato signal from the *Lgr5-EGFP/RCL-tdT* organoid-derived epithelial cell monolayers. Time hh:mm. Scale bar: 500 µm.

**Movie S3 (separate file).** Gap closure experiment in + fibroblasts condition. tdTomato signal from the *Lgr5-EGFP/RCL-tdT* organoid-derived epithelial cell monolayers (white) and GFP signal from the fibroblasts (green). Time hh:mm. Scale bar: 500 µm.

**Movie S4 (separate file).** Gap closure experiment in + $PGE_2$ condition. tdTomato signal from the *ROSA26-creERT2;mT/mG* organoid-derived epithelial cell monolayers (white). Time hh:mm. Scale bar: 500 µm.

**Movie S5 (separate file).** Gap closure experiment in + fibroblasts + EP4i condition. tdTomato signal from the *ROSA26-creERT2;mT/mG* organoid-derived epithelial cell monolayers (green) and phase contrast mage for epithelial cells and fibroblasts (grey). Time hh:mm. Scale bar: 500 µm.

**Movie S6 (separate file).** Formation of a hole in the migrating monolayer in + fibroblasts + EP4i condition. tdTomato signal from the *ROSA26-creERT2;mT/mG* organoid-derived epithelial cell monolayers. Time hh:mm. Scale bar: 100 µm.

**Movie S7 (separate file).** Primary intestinal fibroblasts physically interact with the intestinal cysts. Time hh:mm.

**Movie S8 (separate file).** Movement of a crypt-like region during a gap closure experiment in control conditions. tdTomato signal from the *ROSA26-creERT2;mT/mG* organoid-derived epithelial cell monolayers. Time hh:mm. Scale bar: 50 µm.

**Movie S9 (separate file).** Movement of a crypt-like region during a gap closure experiment in +fibroblasts condition. tdTomato signal from the *ROSA26-creERT2;mT/mG* organoid-derived epithelial cell monolayers. Time hh:mm. Scale bar: 50 µm.

**Movie S10 (separate file).** Formation of a hole in the migrating monolayer in control condition. tdTomato signal from the *Lgr5-EGFP/RCL-tdT* organoid-derived epithelial cell monolayers. Time hh:mm. Scale bar: 100 µm.

**Movie S11 (separate file).** Formation of a hole in the migrating monolayer in + fibroblast condition. tdTomato signal from the *Lgr5-EGFP/RCL-tdT* organoid-derived epithelial cell monolayers. Time hh:mm. Scale bar: 100 µm.

**Movie S12 (separate file).** Formation of a hole in the migrating monolayer in + fibroblast conditioned medium condition. tdTomato signal from the *Lgr5-EGFP/RCL-tdT* organoid-derived epithelial cell monolayers. Time hh:mm. Scale bar: 100 µm.

**Movie S13 (separate file).** Gap closure experiment in the empty gap condition. tdTomato signal from the *Lgr5-EGFP/RCL-tdT* organoid-derived epithelial cell monolayers (white) and GFP signal from the fibroblasts (green). Time hh:mm. Scale bar: 500 µm.

**Movie S14 (separate file).** Gap closure experiment in the CDM condition. tdTomato signal from the *ROSA26-creERT2;mT/mG* organoid-derived epithelial cell monolayers (white). Time hh:mm. Scale bar: 500 µm.
